# Transcriptional Profiling of Identified Neurons in Leech

**DOI:** 10.1101/2020.08.17.254631

**Authors:** Elizabeth Heath-Heckman, Shinja Yoo, Christopher Winchell, Maurizio Pellegrino, James Angstadt, Veronica B. Lammardo, Diana Bautista, Francisco F. De-Miguel, David Weisblat

## Abstract

While leeches in the genus *Hirudo* have long been models for neurobiology, the molecular underpinnings of nervous system structure and function in this group remain largely unknown. To begin to bridge this gap, we performed RNASeq on pools of identified neurons of the central nervous system (CNS): sensory T (touch), P (pressure) and N (nociception) neurons; neurosecretory Retzius cells; and ganglia from which these four cell types had been removed. Bioinformatic analyses identified 2,812 putative genes whose expression differed significantly among the samples. These genes clustered into 7 groups which could be associated with one or more of the identified cell types. We verified predicted expression patterns through *in situ* hybridization on whole CNS ganglia, and found that orthologous genes were for the most part similarly expressed in a divergent leech genus, suggesting evolutionarily conserved roles for these genes. Transcriptional profiling allowed us to identify candidate phenotype-defining genes from expanded gene families. Thus, we identified one of eight hyperpolarization-activated cyclic-nucleotide gated (HCN) channels as a candidate for mediating the prominent sag current in P neurons, and found that one of five inositol triphosphate receptors (IP3Rs), representing a sub-family of IP3Rs absent from vertebrate genomes, is expressed with high specificity in T cells. We also identified one of two *piezo* genes, two of ~65 *deg/enac* genes, and one of at least 16 *transient receptor potential* (*trp*) genes as prime candidates for involvement in sensory transduction in the three distinct classes of leech mechanosensory neurons.

## INTRODUCTION

A major evolutionary advantage arising from multicellularity has been the possibility for species to generate “professional” cell types whose highly differentiated and more or less fixed phenotypes allow them to specialize in particular functions. The broad outlines of how this is achieved through cascading interactions of unequal cell divisions and inherited determinants, intercellular signaling, and transcriptional networks is understood, but numerous questions remain. Nowhere is this more evident than in the nervous system, where the diversity of morphologically defined cell types is further complicated by molecular and physiological distinctions (Shekhar et al. 2016; Diamond 2017; Laboissonniere et al. 2017). Large scale transcriptional profiling at the single cell level (scSeq) is a powerful approach to this problem for complex vertebrate nervous systems, but at present this approach suffers from two limitations.

First is the trade-off between sequencing depth and the mRNA content of the starting sample. Low abundance transcripts are apt to be missing from the transcriptional profile altogether for scSeq, and stochastic variation in which transcripts are counted makes it hard to know which profiles mark phenotypically equivalent cells as opposed to subtle but significant sub-types. A second limitation of current scSeq technologies is the need to dissociate the tissue into its constituent cells as part of the procedure. This obviously results in a loss of spatial information regarding cell identity, and uncertainty in how transcriptional profiles correlate with biophysical properties and physiological functions of the profiled cells.

Certain invertebrate nervous systems offer advantages in addressing these problems, just as they proved advantageous for elucidating aspects of neural mechanisms and neural circuits underlying behavior (Sattelle and Buckingham 2006; Selverston 2010; Taghert and Nitabach 2012). For example, their neurons are often larger in size than those in vertebrates--many neurons in gastropod molluscs, for example, are so large that even individual neurons can be transcriptionally profiled to a much greater depth than is possible for mammalian neurons (Katz and Quinlan 2019). In invertebrates, moreover, one or a few similar neurons coordinate functions that in vertebrates require hundreds or thousands of similar neurons. This together with the extensive body of previous work in certain invertebrate systems offers the ability to study individual cells with well-defined physiological properties and functions (Selverston 2010). In addition, the comparative approach inherent in studying a range of invertebrate systems provides an evolutionary perspective to investigations of how neuronal phenotypes are defined at the molecular level.

Among invertebrates, leeches, primarily the medicinal leech species *Hirudo medicinalis* and *H. verbana*, have long been models for neurobiology. Pioneering neuroanatomical studies in the late 19th century (Retzius 1891) laid the basis for work which combines different experimental approaches to study facets of neurobiology and neurodevelopment ranging from behavior to ion channel function in life stages from the embryo to the adult (Kristan et al. 2005; Weisblat 2007; Mladinic et al. 2009; Wagenaar 2015; Del-Bel and De-Miguel 2018; Kuo and Lai 2019).

The leech CNS comprises a ventral nerve cord of 32 segmental ganglia connected at its anterior end to a non-segmental dorsal ganglion. Twenty-one segmental ganglia innervate segments in the midbody of the animal. Anteriorly, four fused segmental ganglia constitute the ventral portion of the head-brain, connected to the non-segmental dorsal ganglion by circumesophageal nerves; seven fused segmental ganglia make up a tail-brain that innervates the posterior sucker.

In *Hirudo*, most segmental ganglia contain approximately 400 bilaterally paired, individually identifiable neurons distributed in a stereotyped manner. Many identified neurons including sensory neurons, motoneurons and the large neuromodulatory serotonergic Retzius neurons are conserved among different segments in each individual, among individuals within each species, and among different species. The physiological characteristics and behavioral roles are well-known for many of these neurons, including three distinct classes of mechanosensory neurons, the T (touch), P (pressure), and N (nociceptive) cells, whose large cell bodies can be visually identified by their size and position within the segmental ganglia (Fig. 1A).

**Fig.1.**
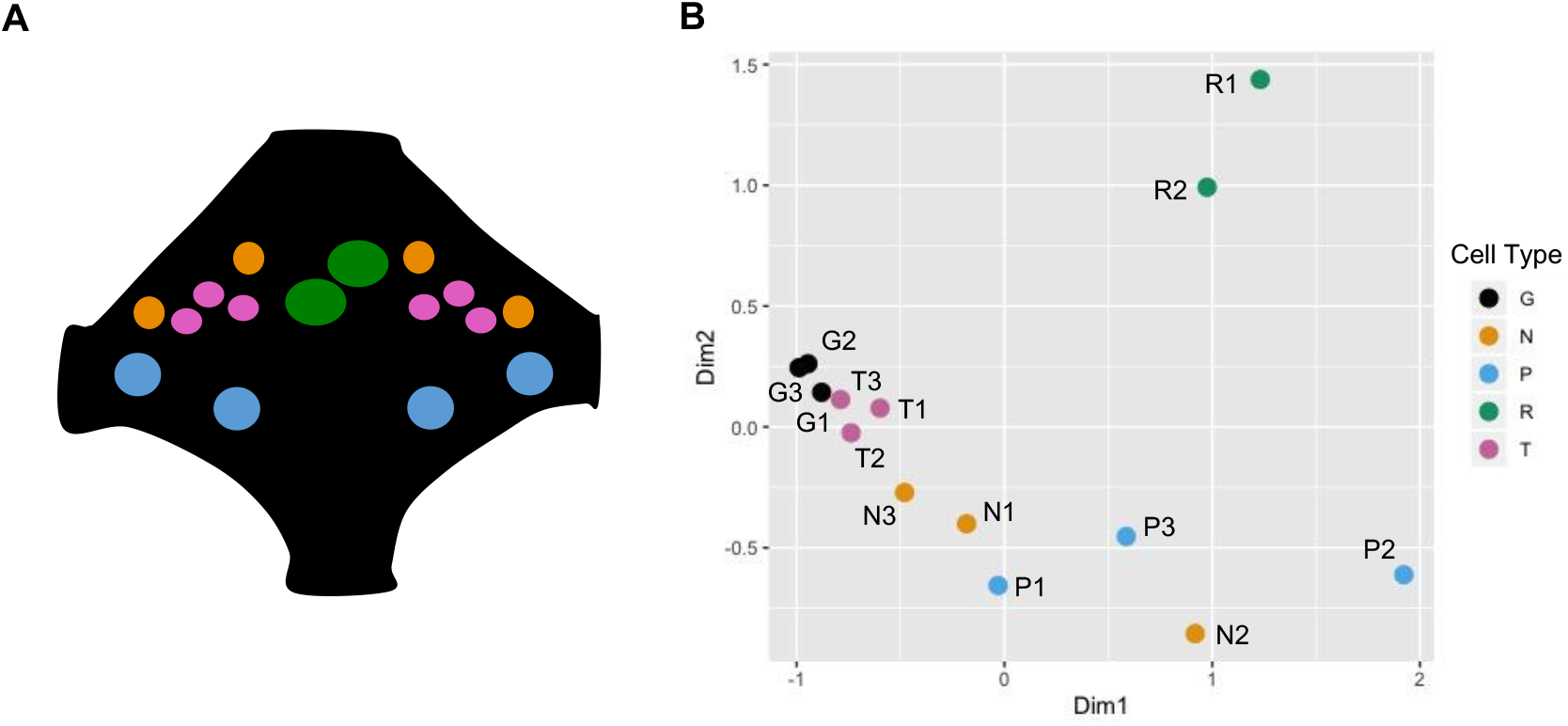
Transcriptional profiles of *H. verbana* neurons and ganglia. A. Schematic of a single *H*. *verbana* segmental ganglion showing the relative positions of the neurons of interest. Each color denotes a sample evaluated in this study: Pink, T neurons (T); Blue, P neurons (P); Orange, N neurons (N), Green, Retzius Neurons (R); Dark Grey/Black, the remainder of the ganglion from which these four cell types had been removed (G). B. A Multi-Dimensional Scaling (MDS) plot showing the relatedness of the transcriptomes of each biological replicate examined in this study.

Three bilateral pairs of T cells exhibit brief action potentials, each followed by a rapidly recovering afterhyperpolarization (Nicholls and Baylor 1968). In response to gentle touch or water flow, T cells fire rapidly adapting bursts of action potentials. The three ipsilateral T cells have partially overlapping dorsal, ventral and lateral receptive fields, respectively, within the ipsilateral body wall (Nicholls and Baylor 1968; Blackshaw 1981). Two bilateral pairs of P cells exhibit somewhat slower action potentials and a marked sag potential in response to hyperpolarizing current injections (Gerard et al. 2012). Their mechanical thresholds are higher than those of T cells, and unlike T cells they give sustained responses during mechanical stimulation (Nicholls and Baylor 1968; Lockery and Sejnowski 1992; Lewis and Kristan 1998; Kretzberg et al. 2016). The medial and lateral cell bodies of P cells innervate partially overlapping dorsal and ventral receptive fields, respectively. P cells exemplify the use of population coding vectors to denote the position of stimuli (Lockery and Sejnowski 1992; Lewis and Kristan 1998). The two pairs of N cells in each ganglion are polymodal nociceptors. In addition to exhibiting the highest threshold to mechanical stimulation of the skin, they respond to other noxious stimuli such as acid, high osmolarity, heat and capsaicin (Pastor et al. 1996). Their action potentials are followed by prominent after-hyperpolarizations (Nicholls and Baylor 1968). In addition to overlapping receptive fields in the body wall, the N cells with more medial cell bodies also innervate the gut (Blackshaw et al. 1982); medial and lateral N cells also differ in their sensitivity to capsaicin and acid (Pastor et al. 1996).

In addition to the three classes of mechanosensory neurons, each ganglion contains a prominent pair of serotonergic neuromodulatory Retzius (Rz) cells (Hagiwara and Morita 1962; Rude et al. 1969). The electrically coupled Retzius cells have the largest cell bodies in the ganglion (Beck et al. 2001); depending on the pattern of electrical activity they may release serotonin from nerve endings, from the axon or from the soma (De-Miguel et al. 2015).

Identified neurons in the adult leech can be removed from ganglia individually; isolated neurons maintain their electrophysiological properties and may grow or form connections with appropriate targets (Chiquet and Nicholls 1987; Nicholls and Hernandez 1989). Thus, the leech *Hirudo* provides a system in which the physiological and behavioral functions of distinct, clearly defined classes of mechanosensory (T, P and N cells) and neurosecretory (Rz) neurons can be examined in detail. Here, we have used the fact that these isolated cells robustly maintain their specific phenotypes to remove and pool neurons of specific phenotypes for RNA extraction and sequencing. For comparison, we have also profiled the transcriptome of the ganglia from which all four of these cell types had been removed. Bioinformatic analyses identified more than two thousand candidate genes whose expression differed significantly among the samples, these genes formed clusters which could be associated to varying extents with one or more of the identified cell types. We verified predicted expression patterns for selected genes through *in situ* hybridization (ISH) on whole leech ganglia. We also found that orthologous genes were (with certain exceptions) similarly expressed in ganglia of a rather distantly related leech *Helobdella austinensis*, suggesting that the genes we assayed play evolutionarily conserved roles in this group. In combination with genome data, transcriptional profiling also allowed us to identify candidate genes for future experiments from among expanded gene families, including specific *piezo*, *deg/enac* and *trp* genes as possible mechanotransducers in the T, P and N neurons.

## RESULTS

### Each neuronal phenotype exhibits a distinctive transcriptional profile

To determine the transcriptional profile of the four cell types, we first created a reference transcriptome by combining the RNA-Seq libraries made from pools of identified T, P, N, and Rz neurons, and libraries made from the remainder of the ganglion after dissection of the four cell types, hereinafter referred to as ganglion-minus (Gm). Three biological replicates were prepared and sequenced for each cell/tissue type, for a total of 15 libraries (Table 1; one of the three Rz replicate libraries yielded a mapping rate 20% lower than any of the other 14 libraries, and was not included in this analysis).

**Table 1.** Sequencing Library and Mapping Information

| Library | Cell Type | Replicate | Number of read pairs | Mapping Rate |
| --- | --- | --- | --- | --- |
| MP53 | P | P1 | 38,227,788 | 84.3 (32231481) |
| MP54 | N | N1 | 21,682,428 | 84.8 (18380876) |
| MP55 | N | N2 | 35,062,153 | 86.6 (30349701) |
| MP56 | T | T1 | 35,013,041 | 82.6 (28904539) |
| MP57 | T | T2 | 36,731,322 | 82.0 (30121670) |
| MP58 | Gm | G1 | 37,025,549 | 80.5 (29794591) |
| MP59 | Gm | G2 | 28,269,748 | 81.0 (22902730) |
| MP60 | Gm | G3 | 32,306,265 | 78.0 (25190050) |
| MP61 | N | N3 | 33,965,761 | 86.3 (29329162) |
| MP62 | T | T3 | 27,553,529 | 84.7 (23332299) |
| MP64 | Retzius | R1 | 34,354,179 | 93.6 (32143423) |
| MP65 | Retzius | R2 | 40,990,667 | 93.2 (38190425) |
| MP66 | P | P2 | 43,127,352 | 90.4 (38967919) |
| MP67 | P | P3 | 41,912,066 | 93.8 (39315980) |

We processed and assembled the resultant reads with Trinity (Grabherr et al. 2011) to obtain a transcriptome containing 113,388 “isoforms”, sequences that may represent variants due to processes such as differential splicing. These isoforms were then grouped into 51,875 “genes”, or unique sequence groups generated by Trinity. The assembled transcriptome has an average sequence length of 786 bp and an N50 of 1173 bp. By comparison, the average transcript length of the 24,432 predicted genes (gene models) in the draft genome assembled for the leech species *Helobdella robusta* is 1.2 kb and their N50 is 1763 bp. Thus, we attribute the discrepancy between the number of “genes” in the *H. verbana* transcriptome and the number of gene models in the *Helobdella* genome to a failure to assemble full *Hirudo* transcripts, so that two or more “genes” correspond to different parts of a single predicted *Helobdella* gene. It is also possible that different potential splice isoforms were separated into separate “genes” when in fact they arise from the same genomic locus. In any case, the sequencing depth achieved by pooling large numbers of phenotypically distinct cell types should prove an important resource for future profiling work at the level of individual neurons.

Our BLAST analysis of the transcriptome assembly revealed that 17,632 (34%) of the “genes” had a significant (e-value < 0.05) BLAST hit in the *H. robusta* gene model database; consistent with the reasoning presented above, there were many cases in which two or more “genes” mapped to a single *Helobdella* gene model. A similar but slightly lower fraction of the *Hirudo* “genes” 15,646 (30%) had a significant BLAST hit against the SwissProt non-redundant database (The UniProt Consortium 2019); we speculate that many of the *Hirudo* “genes” that failed to map to either the *Helobdella* genome or the SwissProt database represent the more rapidly diverging untranslated regions (UTRs) of the transcripts. Finally, when we mapped the original sequence libraries back to the transcriptome we found that all libraries mapped within a range of 78% to 93.6%, suggesting that the transcriptome is representative of the input sequences.

To test the prediction that different cell types have distinct transcriptional profiles, we performed a Multi-Dimensional Scaling (MDS) analysis of the fourteen libraries. As expected, the different sample types segregated into groups roughly corresponding to cell type. The highest degree of internal consistency was for the three T cell transcriptomes and for the three Gm transcriptomes. The Rz cells exhibited the most divergent transcriptional profiles, consistent with their divergent, neurosecretory function (Fig. 1B). In contrast, the replicate P and N cell profiles were interspersed with each other, but distinct from all other samples (Fig. 1B and Discussion). We imagine two possible explanations for the spread in the P and N samples. Given the phenotypic differences between medial and lateral P (and N) cells, (summarized above), one possibility is that medial and lateral P (and N) cells exhibit different transcriptional profiles and are differentially represented among the three replicates. Alternatively, differences in sample collection and processing could give rise to such variation, but why this should only apply to the P and N samples is unclear, since all four cell types were collected in parallel.

### Comparisons among cell types reveal clusters of differentially expressed genes

To determine the functional implications of the differences in transcriptional profiles, we performed all ten possible pairwise comparisons of the five different transcriptomes. These comparisons yielded a set of 2,812 differentially expressed genes (Table 2; see Materials and Methods for details; Supplementary File 1). The total number of genes found in all of the pairwise comparisons summarized in Table 2 was 5,364, reflecting the fact that many of the 2,812 differentially expressed genes occurred in more than one of the pairwise comparisons.

**Table 2.** Numbers of Differentially Enriched Genes in Pairwise Transcriptome Comparisons

|  | Gm | N | P | R | T |
| --- | --- | --- | --- | --- | --- |
| Gm | 0 | 546 | 481 | 1521 | 251 |
| N |  | 0 | 28 | 271 | 171 |
| P |  |  | 0 | 141 | 407 |
| R |  |  |  | 0 | 1547 |
| T |  |  |  |  | 0 |

The similarities and differences in overall transcriptional profiles among the samples, and in the expression of individual genes across samples, were explored using a hierarchical clustering analysis on the biological replicates and on the individual expression profiles of the differentially expressed genes described above (Fig. 2A). Individual samples grouped primarily by cell type, with the exception of two samples (one N and one P) which were also outliers in the MDS plot (Fig. 1B).

**Fig.2.**
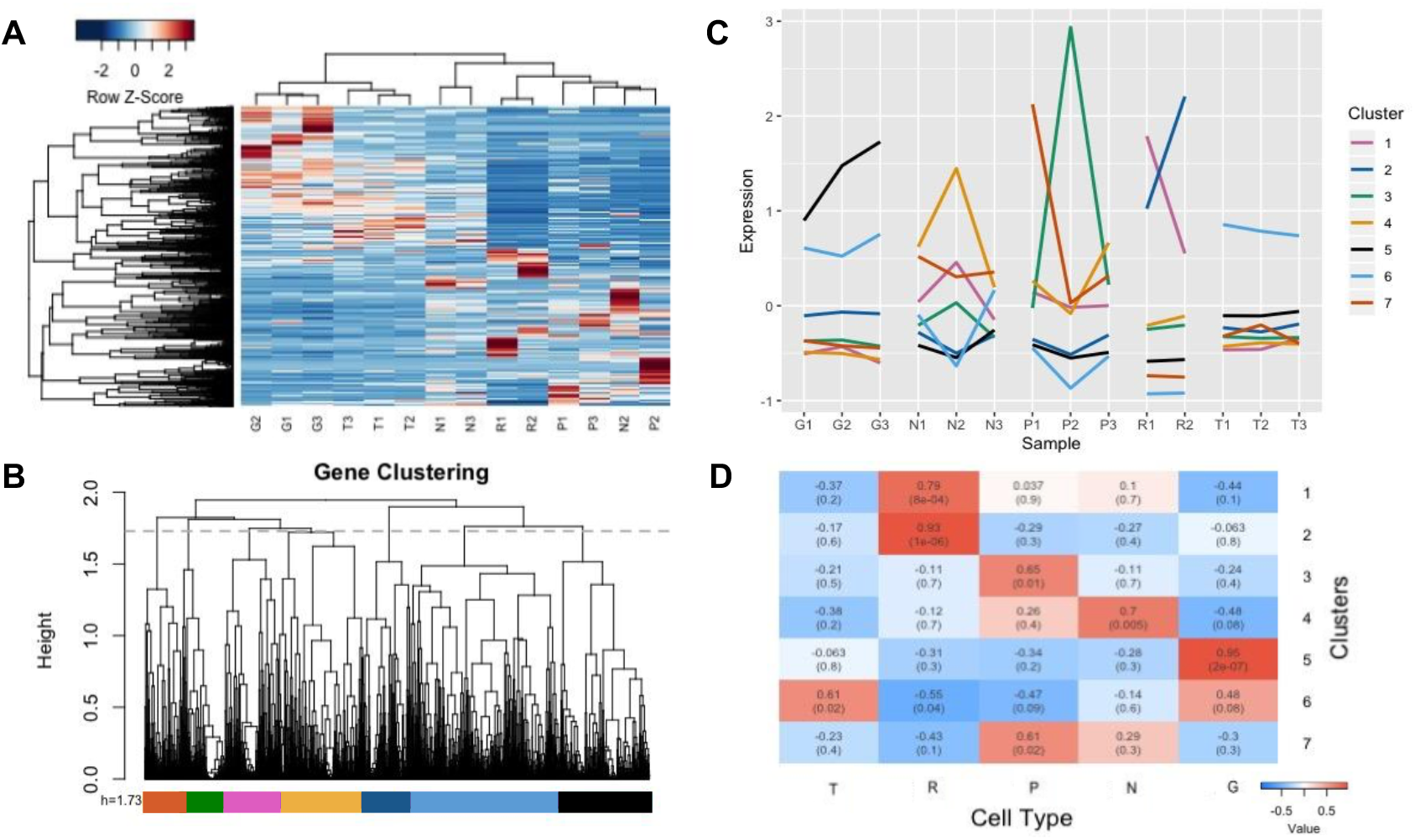
Cluster Analysis and Cellular Enrichment of Differentially Regulated Genes. A. Expression profiles of differentially regulated genes. The heatmap shows the expression patterns of all 2,812 Trinity genes differentially regulated in pairwise analyses of all cell types. Both the biological replicates (X-axis) and differentially regulated genes (Y-axis) have been grouped using hierarchical clustering to reveal patterns of relatedness. B. Parameters chosen for clustering analysis. The dendrogram shown in the Y-axis in (A), with the height cutoff chosen for the following cluster analysis (1.7) shown as a grey dotted line; colored bars denote the resultant clusters. C. Patterns of expression among clusters of differentially regulated genes. The colored lines on the graph indicate the centroid of normalized expression of all genes in the indicated cluster for each biological replicate. Colors denote the same clusters as shown in (B). D. Cluster Enrichment Analysis. The heatmap shows both the correlation coefficient (top) and p-value (bottom, in parentheses) for the cluster-trait analysis showing enrichment of the cluster on the Y-axis in genes expressed in a particular cell type (X-axis).

Naively, one might have expected to obtain five clearly separated clusters of similar gene expression profiles, corresponding to the five sample types. But this was not the case--there was no discrimination height on the gene clustering tree that delineated five clusters correlating with the five sample types (Fig. 2B). At least in retrospect, this initial expectation seems unlikely, given the fact that multiple genes showed up in more than one pairwise comparison (Table 2), and also given the heterogeneity of cell types within the Gm samples. Instead, once the genes were clustered by similarities in expression, we chose a discrimination height on the tree (dotted line in Fig. 2B) that highlighted a workable number of seven clusters of similarly expressed genes for further analysis (Fig. 2B from S1).

The mean expression patterns of the clusters (Fig. 2C and S1) reveal that most of them contain genes that are enriched in particular sample types: genes in Clusters 1 and 2 tended to be enriched in Retzius cells; genes in Cluster 3 tended to be enriched in P cells; genes in Cluster 6 tended to be enriched in T cells and in Gm samples; genes in Cluster 4 tended to be enriched in N cells, and to a lesser extent in P cells; genes in Cluster 5 tended to be enriched in Gm samples; and those in Cluster 7 tended to be enriched in P cells, and to a lesser extent in N cells. We noted, however, that Clusters 1 and 3 in particular seemed to be dominated by a single biological replicate, which calls into question their relevance.

To assess the statistical significance of the correlations between expression of the genes in each cluster with the enrichment in a particular cell type, we performed a cluster-trait analysis on the gene expression clusters (Fig. 2D, see Materials and Methods). Consistent with the neuronal nature of the samples, GO Term enrichment analyses showed that all five samples were dominated by GO Terms for ion channels, receptors and other membrane proteins (data not shown).

### *In situ hybridization* reveals gene expression predicted by transcriptional profiling

To test the validity of the gene expression profiles emerging from the RNASeq analyses, we performed ISH on isolated *Hirudo* ganglia, using probes for genes selected from various clusters (Table 3). This validation was of particular importance because of the possibility for errors in generating the pools of cells used for transcriptional profiling. For example, while the cross-contamination rate for the Rz samples should be near zero, because these cells are unmistakable due to their uniquely large size and position in the ganglion, P cell samples might be contaminated with occasional Leydig cells (Belanger and Orchard 1986), which are similar in size and position to lateral P cells, notwithstanding differences in pigmentation. Similarly, rare cross-contamination of T and N neuron samples may occur because these two cell types, while differing in size, occupy adjacent and variable locations in the anterior portion of the ganglion.

**Table 3.** Genes Used for *in situ* Hybridization Verification of RNASeq Libraries

| Common Name | Cluster | Trinity Gene ID | Helro BLAST (evalue) | SWISS Prot BLAST (evalue) | Average Expression in Each Cell Type in TPM (Transcripts per Kilobase Million) +/- SEM |  |  |  |  |
| --- | --- | --- | --- | --- | --- | --- | --- | --- | --- |
|  |  |  |  |  | T | P | N | Gm | R |
| Aromatic Amino Acid Decarboxylase | 2 | comp24417_c0 | jgi\|Helro1\|186120 (6.91E-70) | P14173.1 (3.12E-6) | 13.7 +/- 2.7 | 8.2 +/- 2.3 | 17.7 +/- 3.1 | 12.5 +/- 2.4 | 2432.5 +/- 465.0 |
| Tryptophan Hydroxylase | 2 | comp20323_c0 | jgi\|Helro1\|79745 (0) | P70080.1 (1.13E-15) | 5.7 +/- 1.0 | 5.5 +/- 1.0 | 5.2 +/- 2.2 | 32.5 +/- 3.1 | 929.0 +/- 28.0 |
| HCN Channel | 4 | comp20462_c0 | jgi\|Helro1\|98055 (8.90E-126) | O88703.1 (10E-101) | 2.8 +/- 1.0 | 31.6 +/- 12.9 | 7.4 +/- 2.0 | 1.3 +/- 0.3 | 0.5 +/- 0.2 |
| Voltage-Gated Potassium Channel | 4 | comp25036_c1 | jgi\|Helro1\|64112 (4.92E-83) | Q9H252 (3.26E-52) | 6.9 +/- 2.0 | 26.1 +/- 11.0 | 5.3 +/- 3.8 | 1.22 +/- 0.3 | 0.0 +/- 0.0 |
| Protocadherin | 4 | comp15454_c0 | jgi\|Helro1\|69346 (1.50E-77) | Q9BZA7.1 (2.58E-48) | 32.6 +/- 13.8 | 511.8 +/- 68.4 | 235.6 +/- 92.2 | 14.3 +/- 1.8 | 7.8 +/- 5.6 |
| Collagen-alpha | 6 | comp13598_c0 | jgi\|Helro1\|110155 (2.52E-17) | Q17RW2.2 (3.39E-11) | 23.5 +/- 5.7 | 0.2 +/- 0.1 | 5.2 +/- 1.0 | 0.6 +/- 0.3 | 0.04 +/- 0.04 |
| Inositol Triphosphate Receptor | 6 | comp24045_c0 | jgi\|Helro1\|162846 (1.84E-69) | P70227.3. (1.44E-06) | 521.5 +/- 186.7 | 22.9 +/- 5.6 | 105.8 +/- 17.4 | 6.9 +/- 1.3 | 3.0 +/- 2.7 |
| Annelid Hypothetical | 7 | comp21991_c0 | jgi\|Helro1\|169916 (6.50E-29) | None | 42.1 +/- 14.8 | 179.6 +/- 80.7 | 103.5 +/- 10.0 | 7.3 +/- 0.7 | 1.0 +/- 0.4 |

For our ISH analysis, we sought to focus on genes with relatively abundant transcripts and relatively selective expression in one of the four cell types under investigation. For this purpose, we first re-examined the set of 2,812 differentially expressed genes to identify those with: 1) BLAST e-values below 0.05 to both the SwissProt and *Helobdella* protein databases; 2) TPM counts above 20 in at least 2 biological replicates, and; 3) a standard error of the mean TPM count not exceeding 40% of the mean of the cell type with the highest expression level. Finally, we excluded genes from Clusters 1 and 3, which were each dominated by a single biological replicate (and thus less likely to represent differentially expressed genes) and Cluster 5, because it represents genes that were *not* expected to be enriched in the neuronal phenotypes of interest, because it is associated with the Gm samples. From the resulting list of 331 genes, we also excluded those with GO Terms indicating mitochondrial or ribosomal functions, leaving a list of 270 candidates (Supplementary File 2) from which we chose genes of interest as described below for ISH analysis (Table 4).

Current ISH protocols for adult *Hirudo* ganglia require the microsurgical removal, prior to fixation, of a protective sheath that encapsulates the ganglion (Coggeshall and Fawcett 1964; Fuchs et al. 1981; Dykes et al. 2004). Unfortunately, this results in the occasional loss or displacement of cell bodies, especially near the lateral edges of the ganglion where the cuts are made. Thus, the counts and spatial distribution of neuronal cell bodies observed in ISH experiments are more variable than in intact ganglia. Nonetheless, as described below, all of the genes tested exhibited characteristic patterns of expression in the ganglion that closely matched the predicted expression from the RNASeq data. In what follows, the gene names used result from molecular phylogenetic and BLAST analyses, as will be explained later in this paper.

Consistent with the serotonergic character of the Rz neurons (e.g., Rude et al. 1969; Henderson 1983), genes encoding proteins involved in biosynthesis and transport of serotonin were prominent components of their transcriptional profile. In particular, Cluster 2 was enriched for transcripts encoding *tryptophan hydroxylase* (*hve-tph*, Trinity Gene ID comp20323_c0) and *dopa decarboxylase (hve-ddc*, comp24417_c0), two enzymes required for serotonin biosynthesis.

*Hve-tph* transcripts were readily detected by ISH in the giant Rz neurons, which occupy a prominent anteromedial location on the ventral surface of the ganglion. A strong ISH signal for *hve-tph* was also observed in two other pairs of smaller serotonergic neurons (cell pairs 21 and 61; Nusbaum and Kristan 1986; Fig. 3A). Surprisingly, however, while *hve-ddc* was readily detected in Rz neurons, we failed to detect an ISH signal for this transcript in cells 21 or 61 (Fig. 3B). The contrast between the ISH results for *hve-tph* and *hve-ddc* was consistent with the differences in the transcript levels obtained from the Gm samples--those samples showed significantly higher counts for *hve-tph* transcripts than did the T, P, or N samples (Fig. 3A), whereas the counts for *hve-ddc* were uniformly low in all but the Rz samples (Fig. 3B).

**Fig. 3.**
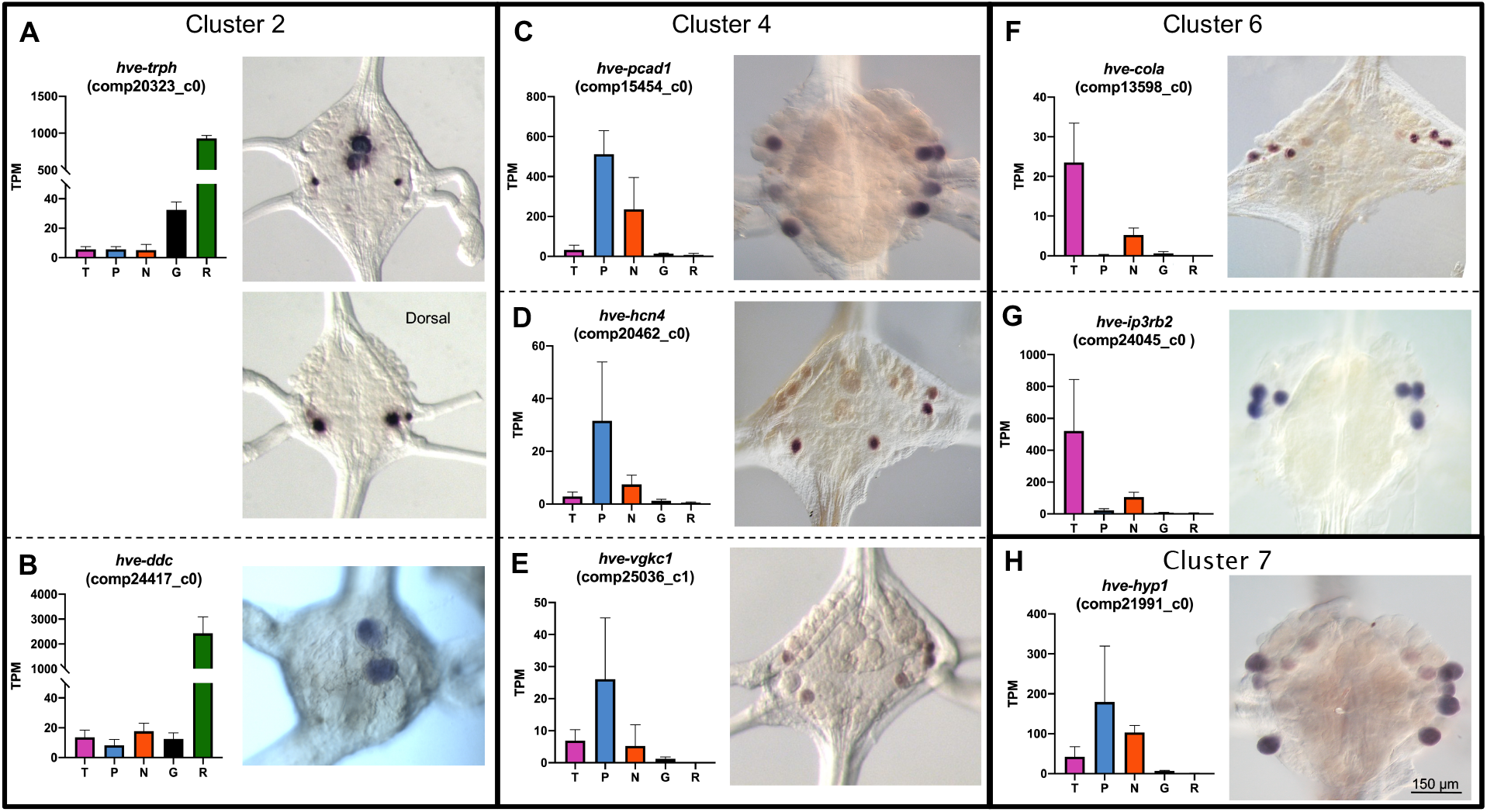
*In situ* hybridization (ISH) verification of expression patterns found by RNASeq. Eight Trinity genes from four clusters were chosen to represent the widest variety of potential staining patterns. A-H. In each panel: the graph at left denotes the expression levels of a Trinity gene in the RNASeq analysis, with the cell type on the X-axis and the average normalized read count of the transcript in transcripts per kilobase million (TPM) on the Y-axis; error bars denote the standard error of the mean; the micrograph at right shows a typical ISH staining pattern for the Trinity gene in an adult *H. verbana* (*Hve*) ganglion. All ganglia are oriented ventral-side up unless otherwise indicated. *tph* = *tryptophan hydroxylase, ddc* = *dopa decarboxylase, pcad1* = *protocadherin 1, hcn4* = *hyperpolarization-activated cyclic nucleotide-gated channel 4, vgkc1* = *voltage-gated potassium channel 1, cola* = *collagen-alpha, ip3rb2* = *inositol triphosphate receptor b2, hyp1* = *hypothetical 1.*

In contrast to the situation for the Rz and T neurons, none of the seven clusters correlated significantly with only the P or N neurons in our analysis, notwithstanding the clear cut phenotypic differences between these neuronal cell types. Rather, most of the transcripts that appeared to be enriched in the transcriptional profile for either the N or P neurons were also present in the other cell type, although at different levels. This is unlikely to be explained by cross-contamination of the samples, because N and P neurons lie in different regions of the ganglia (Fig. 1).

Accordingly, ISH for three genes whose transcripts were relatively abundant and enriched in both the N and P transcriptomes showed expression in both types of neurons (Fig. 3C-3E). These genes, all associated with Cluster 4, encode a protocadherin homolog (*hve-pcad1*, comp15454_c0); a hyperpolarization-activated, cyclic nucleotide-gated cation channel (*hve-hcn4*, comp20462_c0); and a voltage-sensitive potassium channel (*hve-vgkc1*, comp25036_c1). Probes for these three genes labeled combinations of N and P cells that varied somewhat from ganglion to ganglion and even across the midline of individual ganglia (Figure 3C-3E). We attribute this variability, at least in part, to loss or displacement of cells caused by desheathing the ganglia, but an alternative and interesting possibility is that gene expression differs between the medial and lateral members of the N and P cell pairs. Pharmacological and anatomical studies have revealed differences in innervation and sensory coding by the medial and lateral N cells (Pastor et al. 1996), which we expect to reflect differences in their gene expression patterns.

In the transcriptional profiles of T neurons, two genes in Cluster 6 stood out rather unexpectedly, because their biochemical functions seem to correspond to ubiquitously expressed “housekeeping” genes. One encodes a putative collagen-alpha (*hve-cola*, comp13598_c0) and the other encodes a putative receptor for inositol triphosphate (*hve-ip3rb2*, comp24045_c0). For both these genes, the ISH pattern was exceptionally clear, showing three bilateral pairs of labeled neurons in the anterolateral portions of the ganglia correlating with the known positions of the T neuron cell bodies (Fig. 3F and 3G). In light of these results, we speculate that the seemingly increased transcript counts for these genes in the N cell transcriptomes (Fig. 3F and 3G) may represent errors in cell identification during sample preparation.

As a further test for the inferred identity of the neurons expressing *hve-ip3rb2*, and to illustrate the potential for combining molecular and physiological approaches in *Hirudo* ganglia, we used standard techniques to identify T cells by intracellular electrical recordings, and then labeled the identified T neurons by iontophoretic injection of a charged, fixable fluorescent dextran (see Materials and Methods for details). When such preparations were fixed and processed for *hve-ip3rb2* ISH, the ISH product co-localized with the fluorescently labeled neurons, as expected (Fig. 4).

**Fig. 4.**
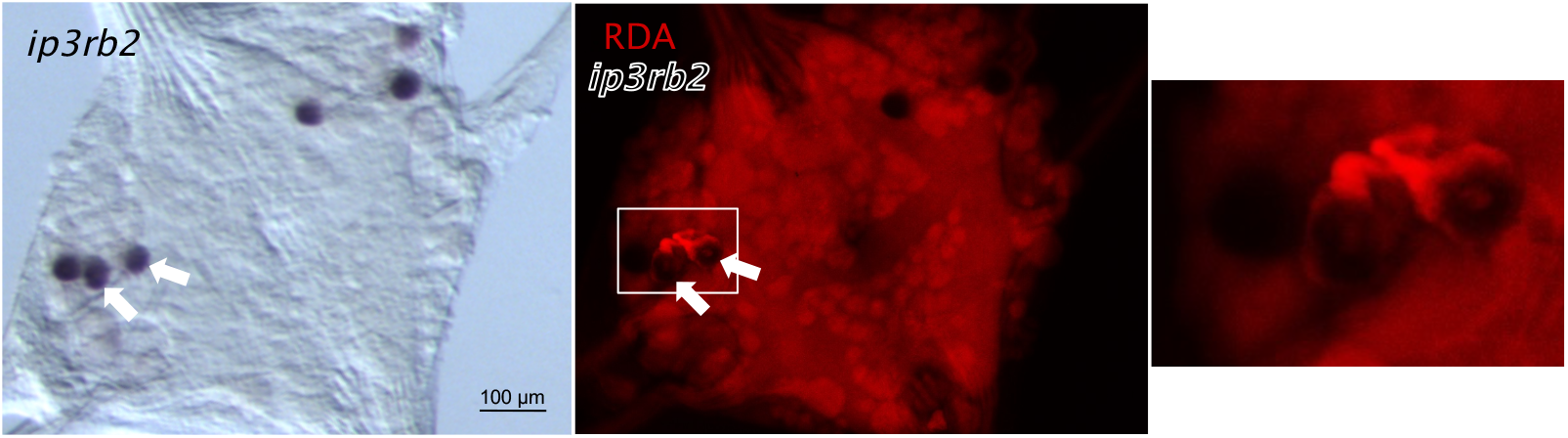
The *ip3rb2* transcript localizes to T neurons. Left panel, brightfield micrograph: ISH for *ip3rb2* shows three bilateral pairs of neurons as expected for the T cells. Center panel, fluorescence micrograph: prior to the *in situ* staining, two of the three T neurons on the left hand side (white arrowheads) had been injected with RDA (red). Right panel, fluorescence micrograph: magnified view of the boxed region in center panel shows that RDA signal in neuronal somata is masked by ISH product. Background signal is due to autofluorescence arising during ISH processing.

In addition to the genes discussed above, we also carried out ISH for a transcript representing a gene for which orthologs are known only from other annelid species. Such genes are candidates for evolutionary novelties, representing hypothetical (hyp) proteins. We chose one such candidate (comp21991_c0, *hve-hyp1)* from Cluster 7. While *hve-hyp1* did not satisfy the criteria for selection above due to a lack of similarity to any proteins in the SWISS-Prot database, we chose it for further analysis to begin to probe how lineage-specific genes may be involved in neuronal specification and function. Consistent with the read counts (Fig. 3H), *hve-hyp1* was expressed primarily in N and P neurons (Fig. 3H).

To explore the extent to which the neuronal markers identified in *Hirudo* may be applicable to other leech species, we identified *Helobdella* orthologs for several of the differentially expressed *Hirudo* genes described above, and then performed ISH for their transcripts on *Helobdella* embryos at stage 10-11 of development, by which time the nervous system is fairly well differentiated and yet ISH can be carried out on intact embryos without dissection (Fig. S2).

As expected, two Rz markers (*hau-tph* and *hau-ddc*) were expressed in the highly conserved serotonergic Rz neurons (Stuart et al. 1987). Intriguingly, *hau-tph* was expressed also in the location of previously described ventrolateral, dorsolateral, and posteromedial serotonergic neurons (Fig. S2; Stuart et al. 1987), but as we had observed for *Hirudo* (Fig. 3), *hau-ddc* was not expressed in these cells.

The expression patterns observed for the *Helobdella* genes *hau-pcad1* and *hau-hcn4* were also similar to those of their *Hirudo* orthologs. Because ISH on *Helobdella* was performed without dissecting the sheath surrounding the ganglion, the expression patterns for these genes were not disrupted by loss or displacement of cells during processing, and clearly repeating patterns were observed. For *hau-pcad1*, four pairs of cells were observed in most ganglia, in positions expected for the bilateral pairs of medial and lateral N and P neurons, but the expression levels were lower in the putative medial N cell than in the other three cells. For *hau-hcn4* three pairs of cells were observed in most ganglia, corresponding to both of the putative P neurons and the lateral but not the medial N neuron.

In contrast to the results for putative Rz, N and P neuron markers, neither of the T cell markers surveyed (*hau-ip3rb2* and *hau-cola)* showed noticeably stronger expression in any particular set of ganglionic neurons (Fig. S2). This result suggests that either the T cells in *Helobdella* use different genes for their specification or function, or that the T cells lag behind other neurons in their development and were not yet expressing these genes at stage 11.

### Molecular phylogeny of amino acid decarboxylases (AADs)

It is paradoxical that the aromatic amino acid decarboxylase gene enriched in Rz neurons was not detected in other known serotonergic neurons in either *Hirudo* or *Helobdella*. One explanation for this observation is that the other neurons are recycling serotonin, taking up serotonin released by the neuromodulatory Rz neurons and then releasing it from their own synapses. This seems unparsimonious, however, given that the non-Rz serotonergic neurons do express tryptophan hydroxylase in both species.

Alternatively, these neurons may use a different aromatic AAD to synthesize serotonin. The *Helobdella* genome encodes four genes annotated as aromatic AADs (AAADs). Three of these genes (JGI gene models 84403, 84539 and 101612) represent comparatively recent duplication events--they are adjacent to one another on genome scaffold 40 and exhibit 57-68% amino acid sequence identity. The fourth gene (JGI gene model 186120), which lies on a separate scaffold and shows only 41-52% sequence identity to the other three, is the ortholog of *hve-ddc*, the gene expressed in Rz neurons.

To explore this issue further, we constructed a molecular phylogeny for the set of AAD sequences obtained by BLASTing a database of non-redundant protein sequences from two model organisms (mouse *Mus musculus* and fruit fly *Drosophila melanogaster*), and three sequenced lophotrochozoan species (*Helobdella robusta*, polychaete annelid *Capitella teleta* and gastropod *Lottia gigantea*). *Mus* and *Drosophila* were chosen to represent the deuterostomes and ecdysozoans, respectively, because the AAADs used for serotonin biosynthesis in these species are known (Scholnick et al. 1986; Juorio et al. 1993).

The genes recovered form two main clades (Fig. 5). One clade comprises acidic amino acid decarboxylases, including a well-supported subclade of glutamic acid decarboxylases (GADs) used to synthesize the neurotransmitter GABA. This GAD subclade included sequences from all five species. The other clade comprises the AAADs. The AAAD clade contains three subclades with broad phylogenetic representation--histidine decarboxylases (HDs, used in histamine biosynthesis, apparently missing in leeches), tyrosine decarboxylases (TDs, used in tyramine and octopamine biosynthesis) and DOPA decarboxylases (DDCs, used in dopamine and serotonin biosynthesis). The gene expressed in leech Rz neurons belongs to the DDC clade, as do the mouse and fly genes used in serotonin biosynthesis. The other three leech AAADs group within the TD subclade. We speculate that one or more of these genes has been co-opted for serotonin biosynthesis in the non-Rz serotonergic neurons but the question remains open. Firstly, the *Hirudo* transcriptome generated here did not contain identifiable orthologs for all three of the *Helobdella* TD genes. Moreover, for the one *Hirudo* transcript (comp21658_c0) that did show sequence similarity to one of the *Helobdella* gene models (101612), the normalized read counts were very low (less than 4 in all samples) in all Gm samples, compared with normalized *hve-tph* read counts of more than 30.

**Fig. 5.**
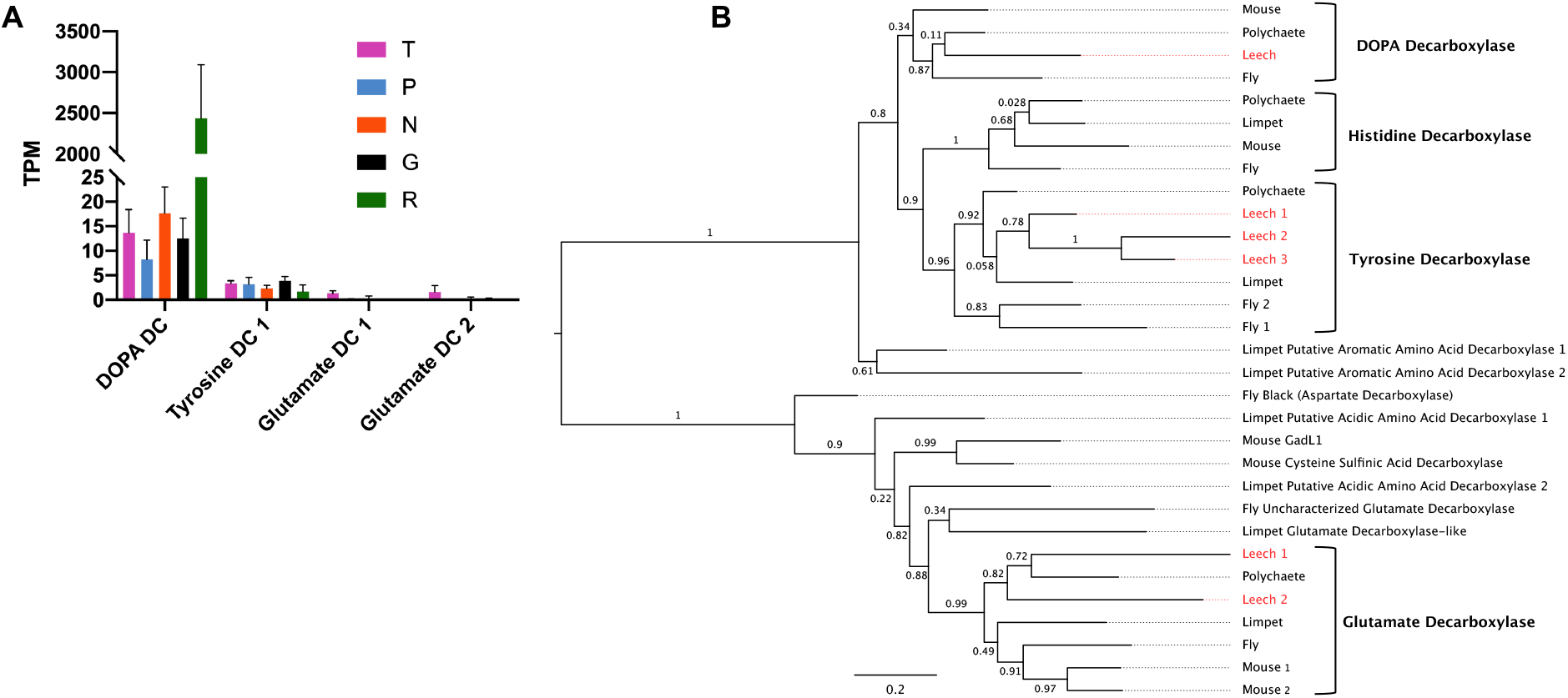
Expression and molecular phylogeny of leech AAD genes A. Expression levels in four cell types and the remainder of the ganglion of the four amino acid decarboxylase genes found in *the Hirudo* transcriptome. Error bars denote the standard error of the mean; TPM = Transcripts per Kilobase Million. B. A maximum-likelihood phylogram including all amino acid decarboxylase genes found in the *Helobdella robusta* genome (in red), only four of which appeared in the *Hirudo* transcriptome. The resolved families of DOPA, Histidine, Tyrosine, and Glutamate Decarboxylases are shown at right. Support values are shown at each node; scale bar = the average number of substitutions per site along each branch. Mouse = *Mus musculus*, Polychaete = *Capitella teleta*, Leech = *Helobdella robusta*, Fly = *Drosophila melanogaster*, Limpet = *Lottia gigantea*

### Expansion of the hcn gene family in leech

One of the prominently upregulated genes in the P and, to a lesser extent, the N cells was a hyperpolarization-activated cyclic nucleotide-gated (HCN) channel (Figures 3 and S2). This channel is a candidate for mediating a prominent, hyperpolarization activated “sag” current that is characteristic of leech P neurons (Baltzley et al. 2010) and heart interneurons (Angstadt and Calabrese 1989). However, evidence of a hyperpolarization-activated current has also been found in the T, P, N, and Rz neurons (Angstadt 1999; Gerard et al. 2012).

In our transcriptome, we found four distinct additional *hcn* transcripts, and examination of another transcriptome (Northcutt et al. 2018) yielded two more, suggesting that *Hirudo* contains at least seven HCN genes. Comparison of these transcripts to the *Helobdella* genome revealed the presence of seven orthologous HCN channel genes and one additional gene as yet found only in *Helobdella*. While this is not a large gene family in absolute terms, it still represents a significant expansion, given that the largest number found in an animal genome to date is four (for mammals; Fig. 6). Our phylogenetic analysis revealed that the HCN gene family has expanded independently in the annelid and vertebrate lineages. Moreover, the expansion of the HCN gene family in annelids appears to be quite recent, as the genome of another annelid (*Capitella teleta*) only encodes one HCN gene, and other lophotrochozoans have at most two (data not shown). Despite their relatively recent emergence, the seven HCN genes in leech appear to have divergent patterns of expression (Figure 6a). Thus, these results exemplify how gene family diversification may contribute to cell phenotype diversification.

**Fig. 6.**
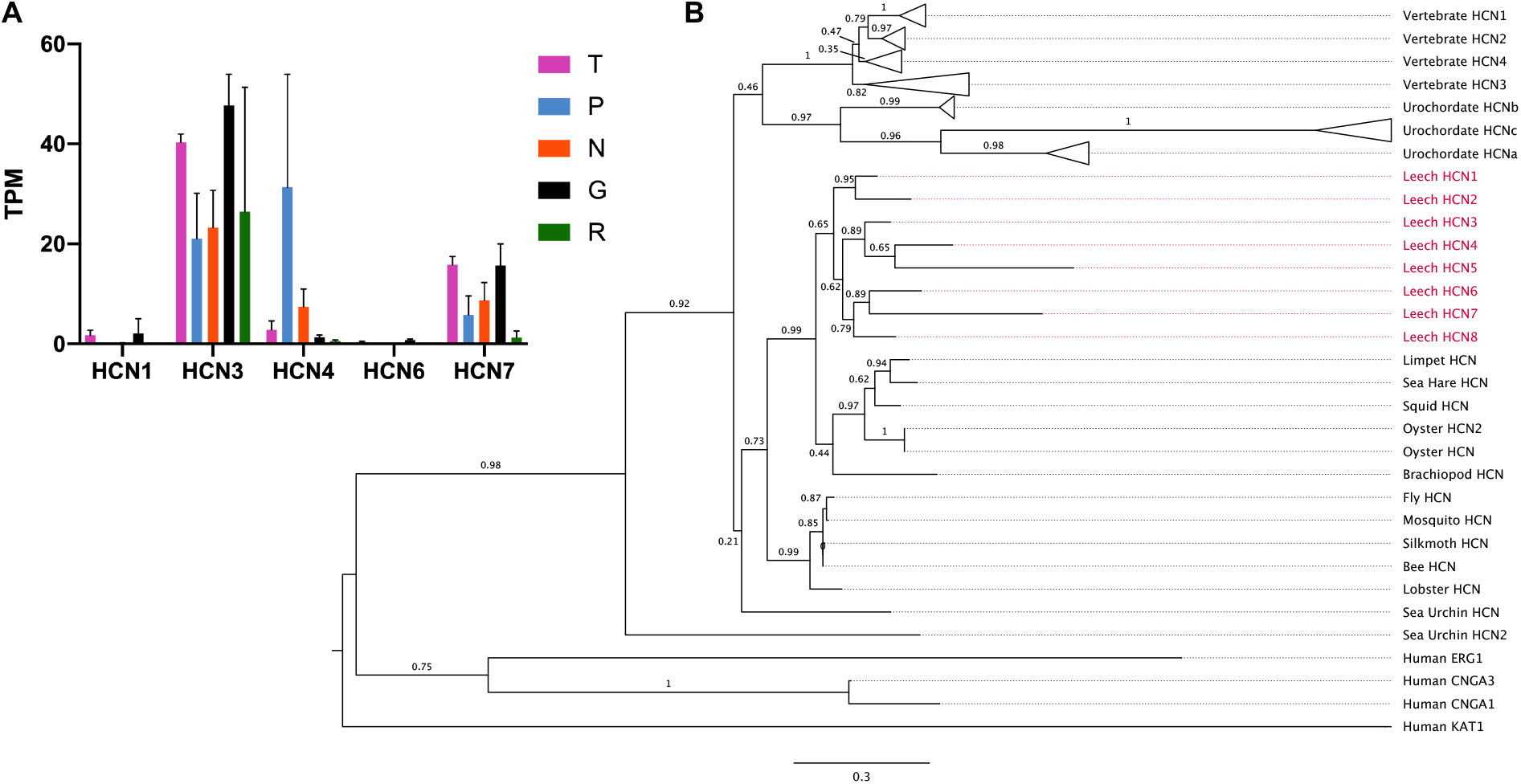
Expression and molecular phylogeny of leech HCN genes. A. Expression levels in four cell types and the remainder of the ganglion of the 5 HCN genes represented in our *Hirudo* transcriptome. Error bars denote the standard error of the mean. TPM = Transcripts per kilobase million. B. A maximum-likelihood phylogeny showing the evolutionary context of the eight HCN genes identified in the *Helobdella robusta* genome (red). Bootstrap values are shown adjacent to each branch; scale bar = the average number of substitutions per site along each branch. Vertebrate = a clade consisting of 11 vertebrate sequences (see figure S3), Urochordate = a clade consisting of *Ciona intestinalis* and *Ciona savignyi*, Leech = *Helobdella robusta*, Limpet = *Lottia gigantea*, Sea Hare = *Aplysia californica*, Squid = *Doryteuthis pealeii*, Oyster = *Crassostrea gigas*, Brachiopod = *Lingula anatina*, Fly = *Drosophila melanogaster*, Mosquito = *Aedes aegypti*, Silkmoth = *Bombyx mori*, Bee = *Apis mellifera*, Lobster = *Panuliris argus*, Sea Urchin = *Strongylocentrotus purpuratus*, Human = *Homo sapiens*

### A phylogenetically distinct IP3 receptor (IP3R) sub-type is preferentially expressed in touch sensitive neurons

Finding an IP3R-encoding transcript among the most prominent elements of the T neuron transcriptional profile, as judged by both relative enrichment and transcript abundance, was unexpected, because we usually think of the IP3Rs as ubiquitously expressed regulators of calcium release from endoplasmic reticulum (Parys and Vervliet 2020). Therefore, having validated this result by ISH (Figs. 3 and 4), we explored the diversity of this gene family in the *Hirudo* transcriptome and the *Helobdella* genome.

Previous work suggests that the bilaterian ancestor had three types of IP3 receptors: IP3RA, IP3RB, which has been lost in the vertebrates, and ryanodine receptors (RyaR; Fig. 7; Alzayady et al. 2015). Our initial BLAST analyses revealed 20 putative IP3R family transcripts in the *Hirudo* transcriptome, compared with five gene loci in the *Helobdella* genome. As IP3Rs are typically large proteins, these five genes were often spread out over two or more (machine annotated) gene models in the *Helobdella* genome. Accordingly, the conceptually translated polypeptides of the 20 *Hirudo ip3r* transcripts map to different regions of four of the five IP3Rs inferred from the *Helobdella* genome, so our data are consistent with a set of at least four mutually orthologous IP3Rs in the two leech species.

**Fig. 7.**
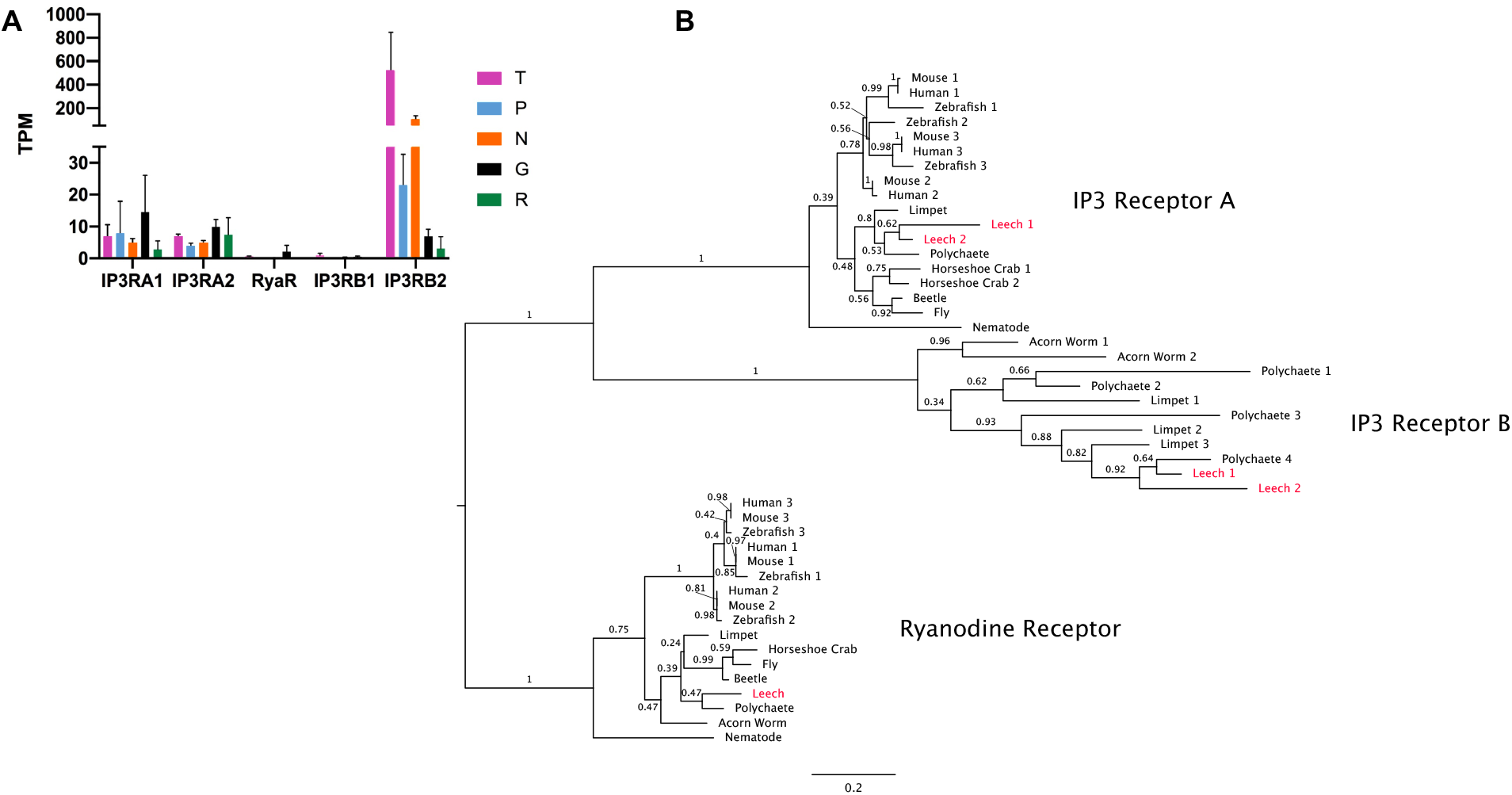
Expression and molecular phylogeny of leech IP3R genes A. Expression levels in four cell types and the remainder of the ganglion of the 5 IP3R transcripts in the *Hirudo* transcriptome. Error bars denote the standard error of the mean; TPM = Transcripts per kilobase million. B. A maximum-likelihood phylogeny including the *Helobdella robusta* orthologs (red) of the transcripts shown in (A), shows their assignment to the three previously identified receptor families: IP3RA, IP3RB and Ryanodine Receptors. Bootstrap values are shown adjacent to each node; scale bar = the average number of substitutions per site along each branch. Mouse = *Mus musculus*, Zebrafish = *Danio rerio*, Human = *Homo sapiens*, Limpet = *Lottia gigantea*, Leech = *Helobdella robusta*, Polychaete = *Capitella teleta*, Horseshoe Crab = *Limulus polyphemus*, Beetle = *Tribolium castaneum*, Fly = *Drosophila melanogaster*, Nematode = *Caenorhabditus elegans*, Acorn Worm = *Saccoglossus kowalevskii*.

The five *Helobdella* IP3Rs include two IP3RA genes, two IP3RB genes, and one RyaR (Fig. 7). Our phylogenetic analysis indicates that the T-enriched IP3R is *ip3rb2.* The IP3RB family of IP3Rs is understudied because it has not been reported in the vertebrates and is also absent in *D. melanogaster* and *C. elegans.* Given the absence of an ISH signal for *ip3rb2* in N neurons, we suspect that the apparently significant levels of *ip3rb2* in the N cell transcriptome may represent contamination of the N cell pools by mis-identified T neurons. In addition to *ip3rb2*, the T cells express *ip3ra1* and *ip3ra2* at levels comparable to the other profiled neurons and the ganglion as a whole.

### Canonical mechanoreceptor candidate genes in the transcriptional profiles of the leech nervous system

The T, P and N neurons in the leech nervous system provide a cellularly well-defined and accessible system in which to study the mechanisms of mechanotransduction in three discrete classes of sensory neurons. As a start toward this goal, we examined the transcriptional profiles for genes related to those that have been implicated in mechanotransduction in other systems, including members of the *piezo*, *trp* and *deg/enac* gene families. This approach is particularly relevant for the *trp* and *enac* gene families, where the large numbers of paralogs would otherwise complicate systematic analysis.

The *piezos* are an ancient gene family with homologs in protozoa, plants and animals (Wu et al. 2017). The *piezos* encode multipass transmembrane proteins that are required for touch sensitivity in mammalian Merkel cells (Woo et al. 2014) and *Drosophila* nociceptors (Kim et al. 2012). In contrast to the Trp and Deg/ENaC channels discussed below, the diverse known physiological roles of Piezos all arise from mechanotransducer functions (Murthy et al. 2017). The *Helobdella* genome contains two *piezo* gene models, but a molecular phylogeny indicates that these are paralogs rather than orthologs of the two mammalian *piezos* (Fig. 8). It was somewhat surprising that no *piezo* gene appeared in the set of differentially expressed genes (Supplementary Table 1). Manual inspection of the transcriptome revealed orthologs of both *Helobdella piezos*, however. Of particular interest, normalized read counts for *Hve-piezo1* appear to be elevated 3-fold in T cells relative to the other neuronal phenotypes. The lack of statistical significance presumably reflects the variability between samples. Normalized read counts for *piezo2* were lower and less variable (Fig. 8). Thus, leech *piezo1* in particular is a candidate for investigation as a mechanotransducer for touch in leech.

**Fig. 8.**
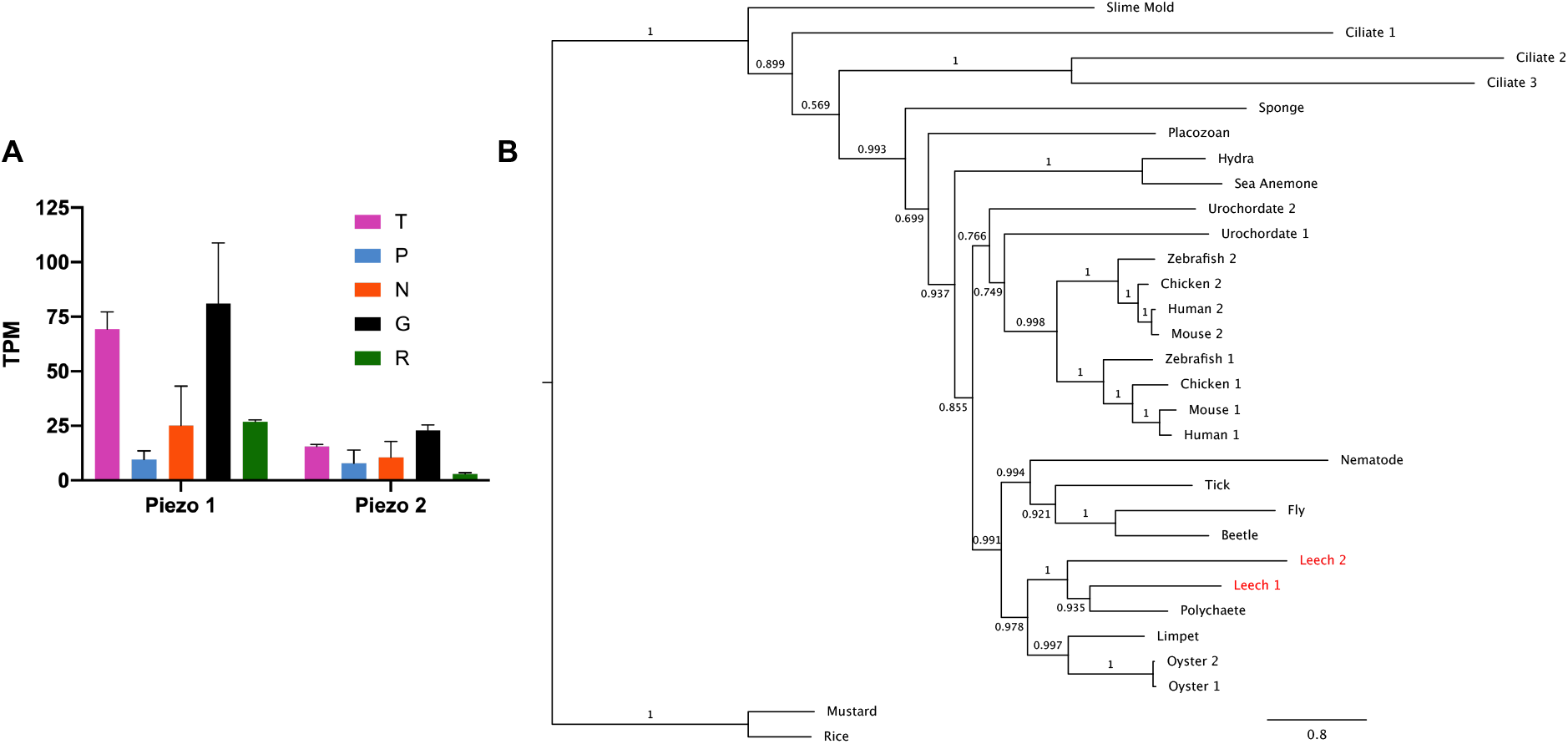
Expression and molecular phylogeny of leech *piezo* genes A. Expression levels in four cell types and the remainder of the ganglion of the two *Hirudo piezo* transcripts. Error bars denote the standard error of the mean. TPM = Transcripts Per Kilobase Million. B. A maximum likelihood phylogeny of Piezo sequences, including the *Helobdella robusta* orthologs (red) of the two *Hirudo* Piezos. Support values are shown at each node; scale bar = the average number of substitutions per site along each branch. Slime Mold = *Dictyostelium discoideum*, Ciliate = *Tetrahymena thermophila*, Sponge = *Amphimedon queenslandica*, Placozoan = *Trichoplax adhaerens*, Hydra = *Hydra vulgaris*, Sea Anemone = *Nematostella vectensis*, Urochordate = *Ciona intestinalis*, Zebrafish = *Danio rerio*, Chicken = *Gallus gallus*, Human = *Homo sapiens*, Mouse = *Mus musculus*, Nematode = *Caenorhabditis elegans*, Tick = *Ixodes scapularis*, Fly = *Drosophila melanogaster*, Beetle = *Tribolium castaneum*, Leech = *Helobdella robusta*, Polychaete = *Capitella teleta*, Limpet = *Lottia gigantea*, Oyster = *Crassostrea gigas*. Mustard = *Arabidopsis thaliana*, Rice = *Oryza brachyantha*.

The *transient receptor potential* (*trp*) genes encode a diverse family of channel proteins, various members of which are implicated in transducing thermal, chemical and mechanical stimuli (Clapham 2003; Nilius and Owsianik 2011; Gees et al. 2012; Julius 2013). Seven sub-families of Trp channels were present in the bilaterian ancestor (Trps A, C, M, ML, P, N, and V) (Peng et al. 2015; Schüler et al. 2015). Many independent *trp* gene duplications have occurred as well. Thus, we find 16 *trp* sequences in the *Hirudo* nervous system transcriptome, each of which has an ortholog in the *Helobdella* genome; these 16 genes include representatives of all sub-families except TrpN and TrpP (Fig. 9). Duplicate genes are present within the TrpC, M and V sub-families; for the M and V sub-families, these appear to represent duplications that had occurred in the bilaterian (M) and protostome (V) ancestors (Peng et al. 2015). Eight genes in the *Hirudo* neuronal transcriptome lie within the *trpA*, *V*, and *M* families, which contain putative sensory genes in other organisms. Among these eight genes, only one, the *Hirudo trpA1* transcript, was enriched in the P cell transcriptome to a statistically significant extent. Thus, this gene is another promising candidate for encoding a sensory transducer in leech.

**Fig. 9.**
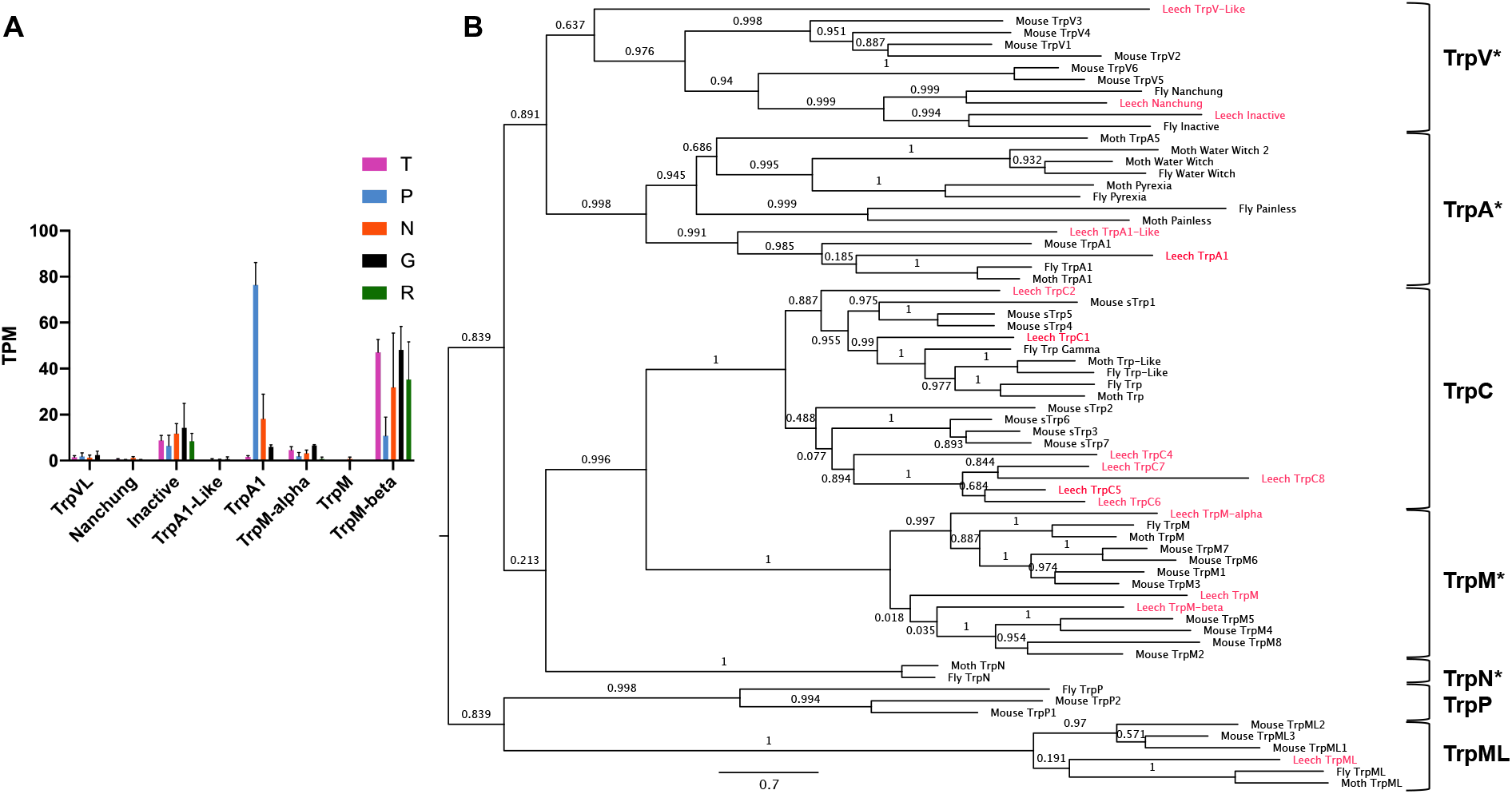
Expression and molecular phylogeny of selected leech TRP genes A. Expression levels in four cell types and in the remainder of the ganglion of the eight *Hirudo* TRP channel transcripts in the TRP V, A, and M families, which are implicated in sensory transduction in other animals. Error bars denote the standard error of the mean; TPM = Transcripts Per Kilobase Million. B. A maximum-likelihood phylogeny of the *Helobdella* orthologs (red) of all 16 TRP channel transcripts found in the *Hirudo* transcriptome, with family groupings labeled at right. Asterisks denote the families implicated in sensory reception in other organisms. Support values are shown at each node; scale bar = the average number of substitutions per site along each branch. Leech = *Helobdella robusta* or *Hirudo verbana* (for TrpA1), Mouse = *Mus musculus*, Fly = *Drosophila melanogaster*, Moth = *Manduca sexta*.

Two Deg/ENaC channels, encoded by the *mec4* and *mec10* genes, mediate mechanotransduction in *C. elegans* touch neurons (Geffeney and Goodman 2012). The *deg/enac* gene family has expanded extensively in the lineage leading to leech from the annelid ancestor; the *Helobdella robusta* genome contains 65 gene models labeled with the Pfam term 008585, corresponding to ENaCs (Simakov et al. 2013). This expansion precludes us from identifying orthologs of *mec4* and *mec10* in leech. Our transcriptome of the *Hirudo* nervous system yielded at least 13 transcripts encoding putative ENaCs, which BLAST to 9 presumptive orthologs in the *Helobdella* genome. Two of these *Hirudo* transcripts appear to be enriched in mechanosensory neurons; one (*Hve-degenac1*, corresponding to the Helobdella gene model 168363) shows moderate read counts (roughly 100 TPM) in the N cell sample and much lower counts (less than 25 TPM) in the other cell types and in the Gm samples. The other (*Hve-degenac2*, corresponding to the *Helobdella* gene model 185250) is expressed at much higher levels in both P and N cell samples (read counts of 1300 and 600 TPM, respectively, Fig. 10).

**Fig. 10.**
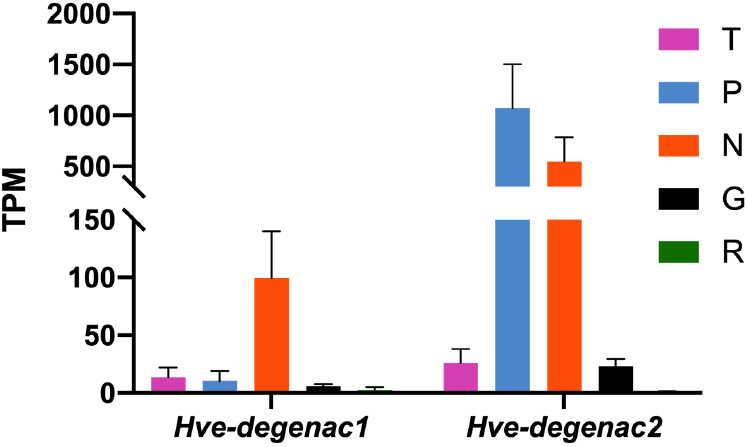
Transcript abundances of differentially regulated Deg/ENaC genes in the *Hirudo* transcriptome. Colored bars denote expression levels in TPM (Transcripts Per Kilobase Million) for each cell type; error bars = the standard error of the mean. *Hve-degenac1* = comp24767_c3, *Hve-degenac2* = comp19322_c0.

## DISCUSSION

### Transcriptional profiles of individually identified, phenotypically distinct neurons

In the work presented here, we have used deep RNA sequencing on pools of individually dissected cells to generate extensive transcriptional profiles for four physiologically and functionally distinct classes of identified neurons from the leech *Hirudo verbana*, as well as for the overall ganglion. Pairwise comparisons between and among the datasets allowed us to generate lists of candidates for genes whose differential expression would contribute to the phenotypic differences among the extensively characterized touch (T), pressure (P), nociceptive (N) and serotonergic neurosecretory Retzius (Rz) neurons of *Hirudo;* in situ hybridization (ISH) allowed us to validate the predicted gene expression patterns of selected genes.

The transcriptomes generated here also set the stage for fine-grained analysis of differences *among* neurons within these various classes, for example by scRNASeq comparisons of medial and lateral N and P neurons within ganglia, and of segment-specific differences in Rz neurons of reproductive segments M5 and M6, versus other midbody segments (Loer et al. 1987; Pastor et al. 1996).

### Expansion of the *hcn* and *ip3r* gene families

Comparative genomic studies indicate that the diversification of bilaterian taxa has been accompanied by lineagespecific expansions of various gene families. Leeches, for example, relative to an inferred annelid ancestor, appear to have undergone an expansion of the gene families encoding innexins, epithelial sodium channels (ENaCs) and homeodomain-containing transcription factors (Kandarian et al. 2012; Simakov et al. 2013), and, as presented here, the hyperpolarization-activated, cyclic nucleotide-gated (HCN) family of ion channels, within the super-family of cyclic nucleotide-gated (CNG) channels.

Finding that one of these *hcn* genes, *hcn4*, is expressed preferentially in P and N cells correlates with previous work showing that P and N cells exhibit enhanced sag voltages and shows how divergence in the regulation of expression among duplicated genes may contribute to divergence in cellular phenotypes. Moreover, this result enables biophysical studies to investigate the extent to which this expansion has been accompanied by functional divergence of the HCN channels.

A second instance of gene family expansion highlighted by this work is in the gene family encoding IP3 receptors (IP3Rs). In the IP3R gene family, three subtypes were inferred for the bilaterian ancestor: IP3RA, IP3RB and RyaR. The leech genome contains duplications of both IP3RA and IP3RB. We found that *ip3rb2* is highly enriched in the T neurons (which respond to light touch) relative to any other cells in the ganglion. Molecular phylogenies indicate that the IP3RB sub-family of IP3 receptors was present in the bilaterian ancestor, but has been lost in the lineage leading to vertebrates. Accordingly, the physiological characterization of these receptors for the most part remain to be determined.

While physiological and pharmacological evidence shows that Rz neurons employ both IP3R- and RyaR-mediated Ca^+2^ release (Trueta et al. 2004; Leon-Pinzon et al. 2014), we detect *ip3ra* and *ip3rb* but not *ryar* transcripts in this cell type. We posit that this is due to a difference in the limit of detection for expression of this transcript, possibly due to sequence artifacts introduced during the *de novo* transcriptome generation, as its expression levels are low in all of our samples (Fig. 5). In any case, this discrepancy highlights the need to complement bioinformatic analyses with direct experimentation.

Given that the T cells express both paralogs of the broadly conserved *ip3ra* sub-family in addition to *ip3rb2*, we speculate that the broadly expressed IP3RAs carry out the housekeeping functions of Ca^+2^ homeostasis, and that the IP3RB family may have been co-opted evolutionarily for T cell-specific functions, the nature of which remain to be determined. We note that mechanical modulation of IP3R-dependent Ca^+2^ release has been observed in mouse endothelium (Wilson et al. 2015).

### Identification of candidate mechanotransducer genes

Transcriptional profiling allows us to sort through dozens of potential candidates and identify specific *piezo, trp and deg/enac* homologs as candidate transduction channels in leech mechanosensory neurons. We identify one of two *piezo* genes, two of ~65 *deg/enac* genes, and one of at least 16 *trp* genes as prime transduction candidates in the three distinct classes of leech mechanosensory neurons. Moreover, these genes appear to be differentially expressed among the three classes of mechanosensory neurons, which correlates with their distinct physiological properties. Specifically, leech *piezo1* appears to be up-regulated in the T neurons, which transduce light touch; this suggests a possible parallel with mammalian *piezo*, which functions to transduce touch in Merkel cells and the associated sensory neurons (Woo et al. 2014). In contrast, the P neurons appear to up-regulate the expression of leech *trpA1* and one of the many leech *deg/enac* genes (*Hve*-*degenac2*, also increased in N cells), while a different *deg/enac* is differentially expressed in the N neurons (*Hve-degenac1*). Since N neurons function as multi-modal nociceptors, responding to salt, acid and heat, we anticipate that other receptors remain to be associated with this class of neurons.

Another intriguing possibility is that the genes that we identify in this study may interact with each other to transduce mechanical signals. For example, in several tissues in vertebrates, Trp channel activation can cause calcium release through IP3 receptors in the ER. While the Trp channels identified in vertebrates, such as TrpM7 (Xiao et al. 2016) and Pkd2 (Delmas 2004), do not have exact orthologs in the leech, the fact that T cells differentially produce a noncanonical IP3 receptor (*ip3rb2*, Fig. 7) and express a member of the TrpM family (*trpM-beta*, Fig. 9) suggests that a similar mechanism may be at play in the light-touch responsive neurons. Given the amenability of the leech for CRISPR/Cas9 and antisense knockdown techniques, we anticipate that future work will be able to test the functional roles of these candidates and other genes in manifesting the distinct phenotypes of the T, P and N neurons.

### Similarities and differences between *Hirudo* and *Helbodella*

Among leeches, the utility of hirudinid species (chiefly *Hirudo verbana* and *H. medicinalis*) as models for physiology and behavior is complemented by the utility of glossiphoniid species (e.g., *Helobdella robusta* and *H. austinensis*) as models for studying early development (Kutschera and Weisblat 2015). In addition to the advantages of being able to apply embryological approaches to the ontogeny of behavior, delineating the similarities and differences among different leech species provides an evolutionary perspective for within this taxon. Comparing the expression of orthologous genes in *Helobdella* and *Hirudo* revealed similarities in expression of *Hau-hcn4, Hau-pcad1, Hau-ddc* and *Hau-tph*, consistent with known similarities in ganglion architecture among leeches (Macagno 1980; Kramer and Kuwada 1983; Kramer and Weisblat 1985; Baltzley et al. 2010).

One curious similarity between *Hirudo* and *Helobdella* is that in both species, the *dopa decarboxylase* (*ddc*) gene that is enriched in serotonergic Rz neurons is not expressed in the smaller, dorsolateral and ventrolateral serotonergic neurons, while *tryptophan hydroxylase* (*tph*) is strongly expressed in all three serotonergic neurons in both species. It remains to be determined whether the decarboxylation step in serotonin biosynthesis is carried out by the product of one of the duplicated genes in the tyrosine decarboxylase clade of AADs in the lateral serotonergic neurons.

A noteworthy difference between the two species is that neither of the *Helobdella* orthologs of the T cell marker genes from *Hirudo*, encoding collagen-alpha and IP3 receptor type B, respectively showed the expected expression patterns in juvenile *Helobdella*. We speculate that the T neurons in the juvenile *Helobdella* used here had not yet differentiated to the point of expressing the *collagen-alpha* or *ip3rb2* genes. An obvious experiment is to test for the expression of these genes in ganglia of adult *Helobdella*, but at present this experiment is technically challenging.

### Conclusions and future directions

In conclusion, the work presented here provides comprehensive transcriptional profiles for four phenotypically distinct classes of identified neurons in the segmental nervous system of the leech *Hirudo verbana*, a tractable model for studying the neurobiological basis of behavior in terms of the properties and connections of individually identified cells. We have used ISH to show that candidate genes exhibit the predicted patterns of expression in *Hirudo* and that orthologous genes are similarly expressed in the nervous system of the leech *Helobdella austinensis*, a tractable model for studying annelid development at the cellular and molecular levels. This lays the basis for future work leveraging the strengths of each system and their underlying similarities to investigate the molecular processes underlying and linking mechanosensation, cell type specification, and behavior. Finally, investigating the differences between the species provides opportunities for studying the evolution of behavioral differences among species at the same level of cellular and molecular detail.

## MATERIALS AND METHODS

### Animals

*Hirudo verbana* (Siddall et al. 2007) were obtained from Leeches USA. *Helobdella austinensis* (Kutschera et al. 2013) were obtained from a laboratory breeding colony at UC Berkeley.

### Isolation of Identified Mechanosensory Neurons

Visually identified T, P, N and Retzius neurons were isolated individually from the central nervous system of adult *H. verbana* as described previously (Dietzel et al. 1986; de-Miguel and Vargas 1997). In brief, leeches were anesthetized by immersion in ice, and short chains of midbody segmental ganglia (with the exception of ganglia 5 and 6) were dissected in Leech Ringer’s solution (Muller and Scott 1981) and pinned in a Sylgard-bottom dish. The ganglia were kept in L-15 culture medium (Gibco) supplemented with 6 mg/ml glucose, 0.1 mg/ml gentamycin (Sigma) and 2% heat-inactivated fetal calf serum (FCS, Microlab). The capsule of each ganglion was opened to expose cell somata and the ganglia chains were incubated for 1 hr in 2 mg/ml collagenase/dispase (Boehringer-Mannheim). After the enzyme treatment, Retzius, T, P and N neurons were identified visually by their size and location in the ganglion. Individual neurons were removed from the ganglia by use of a fire-polished glass pipette. Isolated neurons were rinsed several times in L-15 to sterilize them and remove debris. Three groups of ten leeches were used, yielding three biological replicates, of ~300 cells each, for each cell type. Finally, ganglia from which all the mechanosensory and Retzius neurons had been removed were pooled to create three biological replicates to create the “ganglion” transcriptome.

### RNA Extraction and Sequencing

RNA was isolated from pooled neurons that were stored at −80 in saline solution with 1% (v/v) Recombinant RNasin Ribonuclease Inhibitor (Promega). The RNA was extracted with the Quick-RNA micro prep kit (Zymo) and eluted in 7 ul of nuclease-free water and then made into libraries. The resultant RNA libraries were sequenced by the Functional Genomics Library at UC Berkeley on an Illumina HiSeq 2500 with 2×100 bp reads.

### Transcriptome Assembly and Alignment

We removed sequencing adapters and bases with low quality scores from the reads using Trimmomatic (Bolger et al. 2014) and verified the quality post-trimming with FastQC. If there were overrepresented sequences in the data set postprocessing, we removed them from the data set using custom IlluminaClip parameters in Trimmomatic. To facilitate *de novo* assembly of the transcripts, we performed k-mer normalization using the built-in kmer normalization script in Trinity (Grabherr et al. 2011). In testing phases we also used khmer (Crusoe et al. 2015) with a cutoff value of 20 or 50 and a kmer value of 17 or 20 to compare alternative assembly methods. We then used Trinity (Grabherr et al. 2011) to create *de novo* transcriptomes using these normalized datasets and determined the most representative transcriptome of the datasets. The resultant transcriptome was annotated using BLASTX (Altschul et al. 1990) against the full *Helobdella robusta* genomic protein model database (Simakov et al. 2013) and the SWISSProt Database (The UniProt Consortium 2019). The trimmed reads were then aligned to the transcriptome and the expression levels were determined using Kallisto (Bray et al. 2016).

### Expression Analysis

To identify genes that were differentially expressed among the five sample types, all ten possible pairwise comparisons were performed using the R package edgeR (Robinson et al. 2010) using a p-value of 0.01 after adjusting for multiple comparisons. All differentially regulated genes were then subjected to hierarchical clustering and grouped into 7 clusters using the cutree function and plotted using the R package dendextend (Galili 2015). Cluster-trait correlation analyses were performed using calculations from the R package WGCNA (Langfelder and Horvath 2008). Plots were made using the R packages gplots, ggplot2 (Wickham 2016), and reshape2 (Wickham 2007).

### *In situ* Hybridization

We dissected out ganglia from various midbody segments (M1 through M21, excluding the reproductive segments M5 and M6) of adult *H. verbana* on ice and pinned them onto small, Sylgard-coated plastic petri dishes in Leech Ringer’s solution (Muller and Scott 1981). We then removed the proteinaceous outer sheath using a micro knife (Fine Science Tools) on the ventral face of the ganglion to expose the neuronal cell bodies. At this point, roughly trapezoidal wedges of sylgard to which one or more ganglia were pinned were cut from the dishes and transferred to 1.7 ml polypropylene centrifuge tubes for all further processing. This step simplified the solution changes, reduced the volumes of solution required at each step, reduced damage to the ganglia, and prevented the nerves from folding over the ganglia during the final dehydration steps.

We then fixed the ganglia in 4% paraformaldehyde in 0.5x phosphate-buffered saline (PBS) for 2-3h at room temperature. The ganglia were then brought through a methanol series (5 min at RT in 30%, 50%, 70%, and 90% methanol), washed 3 x 5 min in 100% methanol and stored overnight or until needed in 100% methanol at 4 degrees C, then rehydrated in a methanol series (5 min at RT in 90%, 70%, 50%, and 30% methanol) and washed 2 x 5 min at RT in PBS or PTw (PBS with 0.1% Tween 20). The samples were then digested with 20 ug/mL Proteinase K in PTw for 3-10 min at RT to allow for greater probe access. Digestion was stopped with one 30s wash followed by 2 x 5 min washes in 2 mg/mL glycine in PBS. The ganglia were then washed 3 x 5 min in PBS and post-fixed in 4% paraformaldehyde in PBS for 30-60 min at RT. The fixative was then removed with 2 x 5 min washes in PTw.

To prepare for probe hybridization the ganglia were incubated for 5-10 min in a 1:1 solution of PTw and Hybridization buffer (Hyb: 50% super pure formamide, 5X SSC, 0.2 mg/mL torula RNA, 1X Denhardt’s solution, 0.1 mg/mL heparin, 0.1% Tween-20, 1 mg/mL CHAPS and 9.2 mMol Citric Acid), followed by 5 min in Hyb at RT and then 3-16 hours in fresh Hyb at 65° C. For hybridization, the ganglia were covered with 500-750 ul of Hyb containing 0.2-2.0 ug/mL riboprobe (denatured at 65° C for 25 min prior to addition) and incubated in the probe solution for 16-48 hours at 65° C in a rocking hybridization oven.

To remove unbound probe, the samples were transferred to 2X SSC through a series of 5 min washes at 65° C (Hyb, 1:1 Hyb:SSC, SSC), followed by 20 min 0.2X SSC in PTw, 20 min 0.1X PTw and then 30 sec and 5 min in PTw, all at RT. To reduce nonspecific antibody staining the ganglia were incubated with 1X Western Blocking Reagent (Roche) for 2-3 hours at RT. The samples were then incubated in 1:4000 anti-Digoxigenin conjugated to Alkaline Phosphatase (Roche) in 1X Western Blocking solution for 18 hours at 4° C with rotation. The ganglia were rinsed 3 x 30 sec in PTw, followed by 5 x 1-hour washes in PTw to remove excess antibody.

To visualize the antibody staining, the ganglia, PTw was replaced with BMPurple reagent (Sigma Aldrich) and incubated at RT in the dark until color was seen. The reaction was terminated by rinsing the samples in PTw 3 x 5 min followed by an hour-long incubation in PTw. The specimens were then cleared in a glycerol series (first 40-50% and then 80%), after which they were unpinned from the sylgard block and mounted for microscopy.

### Dye Tracing

Individual ganglia were removed by dissection, pinned onto triangular-shaped slivers of Sylgard designed to slip into 1.5 ml conical tubes, and then desheathed on the ventral or dorsal side. In some experiments, one or more T cells were injected with fixable fluorescent dye (Dextran, Texas Red 3000 MW Lysine; ThermoFisher; 50 mg/ml in the electrode) by iontophoresis, alternating +5 and −5 nA pulses, each with a duration of 600 ms. Injection times were typically on the order of 10-15 min total. Cells were fixed and processed for *in situ* hybridization as described above.

### Molecular Phylogenetics

All protein sequences were obtained from the NCBI protein database (https://www.ncbi.nlm.nih.gov/protein/), the *H. robusta* genome browser (https://mycocosm.jgi.doe.gov/Helro1/Helro1.home.html), the *L. gigantea* genome browser (https://mycocosm.jgi.doe.gov/Lotgi1/Lotgi1.home.html), or the *C. teleta* genome browser (https://mycocosm.jgi.doe.gov/Capca1/Capca1.home.html). Protein alignments were generated via the phylogeny.fr website (Dereeper et al. 2008) using MUSCLE (Edgar 2004) and curated using Gblocks or Noisy curation alignment software (Castresana 2000; Dress et al. 2008). Finally, the maximum-likelihood phylogenies were generated with PhyML (Guindon et al. 2010) using 100 replicates during the bootstrapping process. All sequences used in these analyses can be found in Supplementary File 3.

## DATA AVAILABILITY STATEMENT

The data underlying this article are available in the NCBI BioProject Database at https://www.ncbi.nlm.nih.gov/bioproject/, and can be accessed under the ID PRJNA656790.

## ACKNOWLEDGEMENTS

We thank Lidia Szczupak and Krista Todd for helpful comments and discussions. This work was supported by: the University of California Institute for Mexico and the United States (Grant 2012 to D.A.W., D.B. and FFdM); the Human Frontier Science Program (grant RGP0060/2019 to D.A.W. and F.F.dM. and others), and the National Institutes of Health (NRSA F32NS095665 to E.H-H).

## REFERENCES

Altschul SF, Gish W, Miller W, Myers EW, Lipman DJ. 1990. Basic local alignment search tool. J Mol Biol. 215(3):403–410.

Alzayady KJ, Sebé-Pedrós A, Chandrasekhar R, Wang L, Ruiz-Trillo I, Yule DI. 2015. Tracing the Evolutionary History of Inositol, 1, 4, 5-Trisphosphate Receptor: Insights from Analyses of Capsaspora owczarzaki Ca2+ Release Channel Orthologs. Mol Biol Evol. 32(9):2236–2253.

Angstadt JD. 1999. Persistent inward currents in cultured Retzius cells of the medicinal leech. J Comp Physiol [A]. 184(1):49–61.

Angstadt JD, Calabrese RL. 1989. A hyperpolarization-activated inward current in heart interneurons of the medicinal leech. J Neurosci Off J Soc Neurosci. 9(8):2846–2857.

Baltzley MJ, Gaudry Q, Kristan WB. 2010. Species-specific behavioral patterns correlate with differences in synaptic connections between homologous mechanosensory neurons. J Comp Physiol A. 196(3):181–197.

Beck A, Lohr C, Nett W, Deitmer JW. 2001. Bursting activity in leech Retzius neurons induced by low external chloride. Pflugers Arch. 442(2):263–272.

Belanger JH, Orchard I. 1986. Leydig cells: octopaminergic neurons in the leech. Brain Res. 382(2):387–391.

Blackshaw SE. 1981. Morphology and distribution of touch cell terminals in the skin of the leech. J Physiol. 320(1):219–228.

Blackshaw SE, Nicholls JG, Parnas I. 1982. Physiological responses, receptive fields and terminal arborizations of nociceptive cells in the leech. J Physiol. 326(1):251–260.

Bolger AM, Lohse M, Usadel B. 2014. Trimmomatic: a flexible trimmer for Illumina sequence data. Bioinformatics. 30(15):2114–2120.

Bray NL, Pimentel H, Melsted P, Pachter L. 2016. Near-optimal probabilistic RNA-seq quantification. Nat Biotechnol. 34(5):525–527.

Castresana J. 2000. Selection of Conserved Blocks from Multiple Alignments for Their Use in Phylogenetic Analysis. Mol Biol Evol. 17(4):540–552.

Chiquet M, Nicholls JG. 1987. Neurite outgrowth and synapse formation by identified leech neurones in culture. J Exp Biol. 132(1):191–206.

Clapham DE. 2003. TRP channels as cellular sensors. Nature. 426(6966):517–524.

Coggeshall RE, Fawcett DW. 1964. The fine structure of the central nervous system of the leech, *Hirudo medicinalis*. J Neurophysiol. 27:229–289.

Crusoe MR, Alameldin HF, Awad S, Boucher E, Caldwell A, Cartwright R, Charbonneau A, Constantinides B, Edvenson G, Fay S, et al. 2015. The khmer software package: enabling efficient nucleotide sequence analysis. F1000Research. 4:900.

Del-Bel E, De-Miguel FF. 2018. Extrasynaptic Neurotransmission Mediated by Exocytosis and Diffusive Release of Transmitter Substances. Front Synaptic Neurosci. 10.

Delmas P. 2004. Polycystins: From Mechanosensation to Gene Regulation. Cell. 118(2):145–148.

De-Miguel FF, Leon-Pinzon C, Noguez P, Mendez B. 2015. Serotonin release from the neuronal cell body and its long-lasting effects on the nervous system. Philos Trans R Soc B Biol Sci. 370(1672):20140196.

Dereeper A, Guignon V, Blanc G, Audic S, Buffet S, Chevenet F, Dufayard J-F, Guindon S, Lefort V, Lescot M, et al. 2008. Phylogeny.fr: robust phylogenetic analysis for the non-specialist. Nucleic Acids Res. 36:W465–469.

Diamond JS. 2017. Inhibitory Interneurons in the Retina: Types, Circuitry, and Function. Annu Rev Vis Sci. 3:1–24.

Dietzel ID, Drapeau P, Nicholls JG. 1986. Voltage dependence of 5-hydroxytryptamine release at a synapse between identified leech neurones in culture. J Physiol. 372(1):191–205.

Dress AW, Flamm C, Fritzsch G, Grünewald S, Kruspe M, Prohaska SJ, Stadler PF. 2008. Noisy: Identification of problematic columns in multiple sequence alignments. Algorithms Mol Biol AMB. 3:7.

Dykes IM, Freeman FM, Bacon JP, Davies JA. 2004. Molecular basis of gap junctional communication in the CNS of the leech Hirudo medicinalis. J Neurosci Off J Soc Neurosci. 24(4): 886–894.

Edgar RC. 2004. MUSCLE: multiple sequence alignment with high accuracy and high throughput. Nucleic Acids Res. 32(5):1792–1797.

Fuchs PA, Nicholls JG, Ready DF. 1981. Membrane properties and selective connexions of identified leech neurones in culture. J Physiol. 316(1):203–223.

Galili T. 2015. dendextend: an R package for visualizing, adjusting and comparing trees of hierarchical clustering. Bioinforma Oxf Engl. 31(22):3718–3720.

Gees M, Owsianik G, Nilius B, Voets T. 2012. TRP channels. Compr Physiol. 2(1):563–608.

Geffeney SL, Goodman MB. 2012. How we feel: ion channel partnerships that detect mechanical inputs and give rise to touch and pain perception. Neuron. 74(4):609–619.

Gerard E, Hochstrate P, Dierkes P-W, Coulon P. 2012. Functional properties and cell type specific distribution of Ih channels in leech neurons. J Exp Biol. 215(2):227–238.

Grabherr MG, Haas BJ, Yassour M, Levin JZ, Thompson DA, Amit I, Adiconis X, Fan L, Raychowdhury R, Zeng Q, et al. 2011. Trinity: reconstructing a full-length transcriptome without a genome from RNA-Seq data. Nat Biotechnol. 29(7):644–652.

Guindon S, Dufayard J-F, Lefort V, Anisimova M, Hordijk W, Gascuel O. 2010. New Algorithms and Methods to Estimate Maximum-Likelihood Phylogenies: Assessing the Performance of PhyML 3.0. Syst Biol. 59(3):307–321.

Gustav Retzius. 1891. Zur Kenntniss des centralen Nervensystems der Würmer. In: Biologische Untersuchungen, Neue Folge II. Samson & Wallin. p. 1–28.

Hagiwara S, Morita H. 1962. Electrotonic transmission between two nerve cells in leech ganglion. J Neurophysiol. 25:721–731.

Henderson LP. 1983. The role of 5-hydroxytryptamine as a transmitter between identified leech neurones in culture. J Physiol. 339(1):309–324.

Julius D. 2013. TRP Channels and Pain. Annu Rev Cell Dev Biol. 29(1):355–384.

Juorio AV, Li XM, Walz W, Paterson IA. 1993. Decarboxylation of L-dopa by cultured mouse astrocytes. Brain Res. 626(1–2):306–309.

Kandarian B, Sethi J, Wu A, Baker M, Yazdani N, Kym E, Sanchez A, Edsall L, Gaasterland T, Macagno E. 2012. The medicinal leech genome encodes 21 innexin genes: different combinations are expressed by identified central neurons. Dev Genes Evol. 222(1):29–44.

Kim SE, Coste B, Chadha A, Cook B, Patapoutian A. 2012. The role of Drosophila Piezo in mechanical nociception. Nature. 483(7388):209–212.

Kramer AP, Kuwada JY. 1983. Formation of the receptive fields of leech mechanosensory neurons during embryonic development. J Neurosci Off J Soc Neurosci. 3(12):2474–2486.

Kramer AP, Weisblat DA. 1985. Developmental neural kinship groups in the leech. J Neurosci. 5(2):388–407.

Kretzberg J, Pirschel F, Fathiazar E, Hilgen G. 2016. Encoding of Tactile Stimuli by Mechanoreceptors and Interneurons of the Medicinal Leech. Front Physiol. 7:506.

Kristan WB, Calabrese RL, Friesen WO. 2005. Neuronal control of leech behavior. Prog Neurobiol. 76(5):279–327.

Kuo D-H, Lai Y-T. 2019. On the origin of leeches by evolution of development. Dev Growth Differ. 61(1):43–57.

Kutschera U, Langguth H, Kuo D-H, Weisblat DA, Shankland M. 2013. Description of a new leech species from North America, Helobdella austinensis n. sp. (Hirudinea: Glossiphoniidae), with observations on its feeding behaviour. Zoosystematics Evol. 89(2):239–246.

Kutschera U, Weisblat DA. 2015. Leeches of the genus Helobdella as model organisms for Evo-Devo studies. Theory Biosci Theor Den Biowissenschaften. 134(3–4):93–104.

Laboissonniere LA, Sonoda T, Lee SK, Trimarchi JM, Schmidt TM. 2017. Single-cell RNA-Seq of Defined Subsets of Retinal Ganglion Cells. JoVE J Vis Exp. (123):e55229.

Langfelder P, Horvath S. 2008. WGCNA: an R package for weighted correlation network analysis. BMC Bioinformatics. 9(1):559.

Leon-Pinzon C, Cercós MG, Noguez P, Trueta C, De-Miguel FF. 2014. Exocytosis of serotonin from the neuronal soma is sustained by a serotonin and calcium-dependent feedback loop. Front Cell Neurosci. 8.

Lewis JE, Kristan WB. 1998. A neuronal network for computing population vectors in the leech. Nature. 391(6662):76–79.

Lockery SR, Sejnowski TJ. 1992. Distributed processing of sensory information in the leech. III. A dynamical neural network model of the local bending reflex. J Neurosci. 12(10):3877–3895.

Loer C, Jellies J, Kristan W. 1987. Segment-specific morphogenesis of leech Retzius neurons requires particular peripheral targets. J Neurosci. 7(9):2630–2638.

Macagno ER. 1980. Number and distribution of neurons in leech segmental ganglia. J Comp Neurol. 190(2):283–302.

de-Miguel FE, Vargas J. 1997. Different determinants on growth and synapse formation in cultured neurons. Neuroreport. 8(3):761–765.

Mladinic M, Muller KJ, Nicholls JG. 2009. Central nervous system regeneration: from leech to opossum: Central nervous system regeneration. J Physiol. 587(12):2775–2782.

Muller KJ, Scott SA. 1981. Transmission at a ‘direct’ electrical connexion mediated by an interneurone in the leech. J Physiol. 311(1):565–583.

Murthy SE, Dubin AE, Patapoutian A. 2017. Piezos thrive under pressure: mechanically activated ion channels in health and disease. Nat Rev Mol Cell Biol. 18(12):771–783.

Nicholls JG, Baylor DA. 1968. Specific modalities and receptive fields of sensory neurons in CNS of the leech. J Neurophysiol. 31(5):740–756.

Nicholls JG, Hernandez UG. 1989. Growth and Synapse Formation by Identified Leech Neurones in Culture: A Review. Q J Exp Physiol. 74(6):965–973.

Nilius B, Owsianik G. 2011. The transient receptor potential family of ion channels. Genome Biol. 12(3):218.

Northcutt AJ, Fischer EK, Puhl JG, Mesce KA, Schulz DJ. 2018. An annotated CNS transcriptome of the medicinal leech, Hirudo verbana: De novo sequencing to characterize genes associated with nervous system activity. PloS One. 13(7):e0201206.

Nusbaum MP, Kristan WB. 1986. Swim initiation in the leech by serotonin-containing interneurones, cells 21 and 61. J Exp Biol. 122:277–302.

Parys JB, Vervliet T. 2020. New Insights in the IP3 Receptor and Its Regulation. Adv Exp Med Biol. 1131:243–270.

Pastor J, Soria B, Belmonte C. 1996. Properties of the nociceptive neurons of the leech segmental ganglion. J Neurophysiol. 75(6):2268–2279.

Peng G, Shi X, Kadowaki T. 2015. Evolution of TRP channels inferred by their classification in diverse animal species. Mol Phylogenet Evol. 84:145–157.

Robinson MD, McCarthy DJ, Smyth GK. 2010. edgeR: a Bioconductor package for differential expression analysis of digital gene expression data. Bioinformatics. 26(1):139–140.

Rude S, Coggeshall E, Van Orden LS. 1969. Chemical and ultrastructural identification of 5-hydroxytryptamine in an identified neuron. J Cell Biol. 41(3):832–854.

Sattelle DB, Buckingham SD. 2006. Invertebrate studies and their ongoing contributions to neuroscience. Invertebr Neurosci IN. 6(1):1–3.

Scholnick SB, Bray SJ, Morgan BA, McCormick CA, Hirsh J. 1986. CNS and hypoderm regulatory elements of the Drosophila melanogaster dopa decarboxylase gene. Science. 234(4779):998–1002.

Schüler A, Schmitz G, Reft A, Özbek S, Thurm U, Bornberg-Bauer E. 2015. The Rise and Fall of TRP-N, an Ancient Family of Mechanogated Ion Channels, in Metazoa. Genome Biol Evol. 7(6):1713–1727.

Selverston AI. 2010. Invertebrate central pattern generator circuits. Philos Trans R Soc B Biol Sci. 365(1551):2329–2345.

Shekhar K, Lapan SW, Whitney IE, Tran NM, Macosko EZ, Kowalczyk M, Adiconis X, Levin JZ, Nemesh J, Goldman M, et al. 2016. Comprehensive Classification of Retinal Bipolar Neurons by Single-Cell Transcriptomics. Cell. 166(5):1308–323.e30.

Siddall ME, Trontelj P, Utevsky SY, Nkamany M, Macdonald KS. 2007. Diverse molecular data demonstrate that commercially available medicinal leeches are not Hirudo medicinalis. Proc Biol Sci. 274(1617): 1481–1487.

Simakov O, Marletaz F, Cho S-J, Edsinger-Gonzales E, Havlak P, Hellsten U, Kuo D-H, Larsson T, Lv J, Arendt D, et al. 2013. Insights into bilaterian evolution from three spiralian genomes. Nature. 493(7433):526–531.

Stuart DK, Blair SS, Weisblat DA. 1987. Cell lineage, cell death, and the developmental origin of identified serotonin- and dopamine-containing neurons in the leech. J Neurosci. 7(4):1107–1122.

Taghert PH, Nitabach MN. 2012. Peptide neuromodulation in invertebrate model systems. Neuron. 76(1):82–97.

The UniProt Consortium. 2019. UniProt: a worldwide hub of protein knowledge. Nucleic Acids Res. 47(D1):D506–D515.

Trueta C, Sánchez-Armass S, Morales MA, De-Miguel FF. 2004. Calcium-induced calcium release contributes to somatic secretion of serotonin in leech Retzius neurons. J Neurobiol. 61(3):309–316.

Wagenaar DA. 2015. A classic model animal in the 21st century: recent lessons from the leech nervous system. J Exp Biol. 218(21):3353–3359.

Weisblat DA. 2007. Asymmetric Cell Divisions in the Early Embryo of the Leech Helobdella robusta. In: Macieira-Coelho A, editor. Asymmetric Cell Division. Berlin, Heidelberg: Springer. (Progress in Molecular and Subcellular Biology). p. 79–95.

Wickham H. 2007. Reshaping Data with the reshape Package. J Stat Softw. 21(1): 1–20.

Wickham H. 2016. ggplot2: Elegant Graphics for Data Analysis. 2nd ed. Springer International Publishing.

Wilson C, Saunter CD, Girkin JM, McCarron JG. 2015. Pressure-dependent regulation of Ca2+ signalling in the vascular endothelium. J Physiol. 593(24):5231–5253.

Woo S-H, Ranade S, Weyer AD, Dubin AE, Baba Y, Qiu Z, Petrus M, Miyamoto T, Reddy K, Lumpkin EA, et al. 2014. Piezo2 is required for Merkel-cell mechanotransduction. Nature. 509(7502):622–626.

Wu J, Lewis A, Grandl J. 2017. Touch, Tension, and Transduction – the Function and Regulation of Piezo Ion Channels. Trends Biochem Sci. 42(1):57–71.

Xiao E, Chen C, Zhang Y. 2016. The mechanosensor of mesenchymal stem cells: mechanosensitive channel or cytoskeleton? Stem Cell Res Ther. 7.

