## Supplementary Material for "Transcriptional Profiling of Identified Neurons in Leech"

**Supplementary Tables and Figures**

**Table S1: Primers used in this study**

| Common Name | F (*H. verbana*) | R (*H. verbana*) | F (*H. robusta*) | R (*H. robusta*) |
|---|---|---|---|---|
| Inositol Triphosphate Receptor | gagggagaatccgtgcaaata | gaagcactctcaccaacaaatc | ctaggcagtacctatgcatcac | atagcacccatccctcctta |
| Collagen-alpha | ccgtcagttccctggtttc | caagggtgactctggagaaatag | gtattgaaggtcctgtgggaat | agtggactgggaggatcaa |
| Annelid hypothetical | cggcgtgaacatctcatagaa | ctcctcccacaacatggaaa | n/a | n/a |
| Protocadherin | ccagtcacagggtcgaaatag | cccgctaaagtgatcgtgaa | tggctggaatctcaaacttact | cgggcgtcgatgttaaattattc |
| HCN Channel | tttccctgtaatgcctctttga | cggagtccagaggaggataa | cccatgctacaagacttccc | tcgtacgttgcataggataaaca |
| Voltage-Gated Potassium Channel | tactctccgctcaaaagtgc | taatacgactcactataggaagtccttaatgcttgcgc (with t7) | n/a | n/a |
| Aromatic Amino Acid Decarboxylase | gaaggcctttgtctcaggtaa | tgtctgaactcgggacaaataaa | atgatgaagtggctcggtaag | gttgacgtcaccttcgttagt |
| Tryptophan Hydroxylase | tgaccatcctggtttcaaaga | tcgataaaggtttcctgctctactgc | gatggtcacccgtggtttcc | tcgacaagttgaacagtttatcctgg |

**Fig. S1.** Expression patterns of each isolated cluster. Centroids of each cluster are shown in color, while the expression of each Trinity gene belonging to the cluster is shown in light grey.

**Fig. S2.** Expression of neuron-specific transcripts found in *Hirudo* in late-stage *Helobdella* embryos. ISH of stage 11 *Helobdella austinensis* embryos using probes specific to the *Helobdella* orthologs of six of the transcripts verified in figure 3. Probe localization is shown in blue/purple. For each probe, a light micrograph of a chain of 4 mid-body ganglia (left), and a single ganglion at higher magnification (right) are shown. Anterior-posterior (A-P) axis orientation is shown at left.

**Fig. S3.** Full Phylogenetic tree shown in Figure 5, with all members of each clade shown. Scale bar = the average number of substitutions per site along each branch.

Figure S1

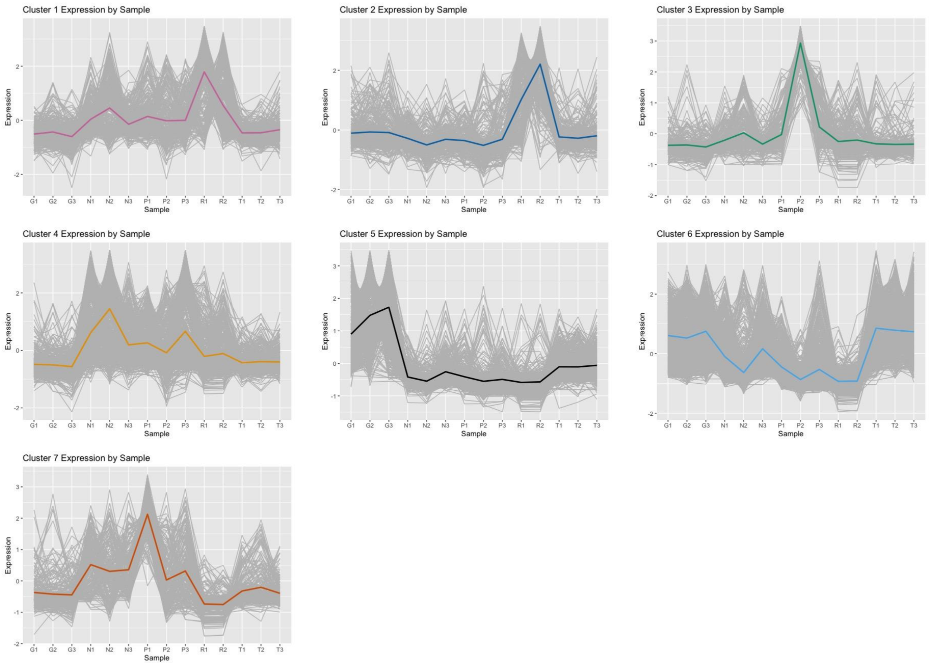


Figure S2

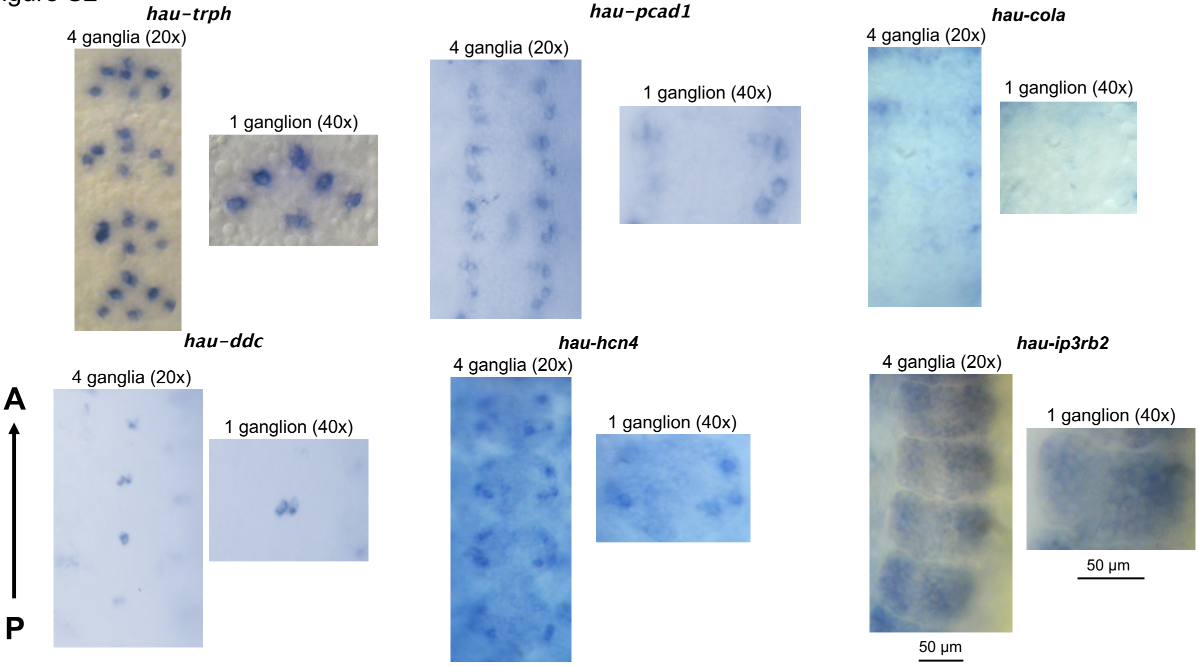

Figure S3

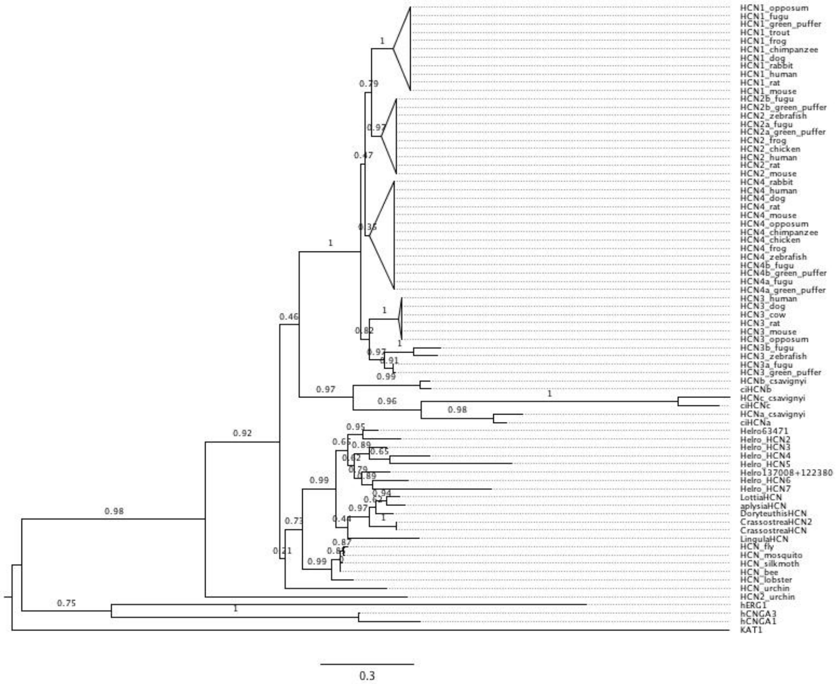
HCN1_opposum
HCN1_fugu
HCN1_green_puffer
HCN1_trout
HCN1_frog
HCN1_chimpanzee
HCN1_dog
HCN1_rabbit
HCN1_human
HCN1_rat
HCN1_mouse
HCN2b_fugu
HCN2b_green_puffer
HCN2_zebrafish
HCN2a_fugu
HCN2a_green_puffer
HCN2_frog
HCN2_chicken
HCN2_human
HCN2_rat
HCN2_mouse
HCN4_rabbit
HCN4_human
HCN4_dog
HCN4_rat
HCN4_mouse
HCN4_opposum
HCN4_chimpanzee
HCN4_chicken
HCN4_frog
HCN4_zebrafish
HCN4b_fugu
HCN4b_green_puffer
HCN4a_fugu
HCN4a_green_puffer
HCN3_human
HCN3_dog
HCN3_cow
HCN3_rat
HCN3_mouse
HCN3_opposum
HCN3b_fugu
HCN3_zebrafish
HCN3a_fugu
HCN3_green_puffer
HCNb_csavignyi
ciHCNb
HCNc_csavignyi
ciHCNc
HCNa_csavignyi
ciHCNa
Helro63471
Helro_HCN2
Helro_HCN3
Helro_HCN4
Helro_HCN5
Helro137008+122380
Helro_HCN6
Helro_HCN7
LottiaHCN
aplysiaHCN
DoryteuthisHCN
CrassostreaHCN2
CrassostreaHCN
LingulaHCN
HCN_fly
HCN_mosquito
HCN_silkmoth
HCN_bee
HCN_lobster
HCN_urchin
HCN2_urchin
hERG1
hCNGA3
hCNGA1
KAT1
0.3
